# Circulating immune cell phenotype dynamics reflect the strength of tumor-immune cell interactions in patients during immunotherapy

**DOI:** 10.1101/2020.03.24.993923

**Authors:** Jason I Griffiths, Pierre Wallet, Lance T. Pflieger, David Stenehjem, Xuan Liu, Patrick A. Cosgrove, Neena A. Leggett, Jasmine McQuerry, Gajendra Shrestha, Maura Rossetti, Gemalene Sunga, Philip J. Moos, Frederick R. Adler, Jeffrey T. Chang, Sunil Sharma, Andrea H. Bild

## Abstract

The extent that immune cell phenotypes in the peripheral blood reflect within-tumor immune activity prior to and early in cancer therapy is unclear. To address this question, we studied the population dynamics of tumor and immune cells, and immune phenotypic changes, using clinical tumor and immune cell measurements and single cell genomic analyses. These samples were serially obtained from a cohort of advanced gastrointestinal cancer patients enrolled on a trial with chemotherapy and immunotherapy. Using an ecological population model, fitted to clinical tumor burden and immune cell abundance data from each patient, we find evidence of a strong tumor-circulating immune cell interaction in responder patients, but not those patients that progress on treatment. Upon initiation of therapy, immune cell abundance increased rapidly in responsive patients, and once the peak level is reached, tumor burden decreases, similar to models of predator-prey interactions; these dynamic patterns were absent in non-responder patients. To interrogate phenotype dynamics of circulating immune cells, we performed single cell RNA sequencing at serial time points during treatment. These data show that peripheral immune cell phenotypes were linked to the increased strength of patients’ tumor-immune cell interaction, including increased cytotoxic differentiation and strong activation of interferon signaling in peripheral T-cells in responder patients. Joint modeling of clinical and genomic data highlights the interactions between tumor and immune cell populations and reveals how variation in patient responsiveness can be explained by differences in peripheral immune cell signaling and differentiation soon after the initiation of immunotherapy.

**One sentence summary:** Peripheral immune cell differentiation and signaling, upon initiation of immunotherapy, reflects tumor attacking ability and patient response.

**Significance statement:** The evolution of peripheral immune cell abundance and signaling over time, as well as how these immune cells interact with the tumor, may impact a cancer patient’s response to therapy. By developing an ecological population model, we provide evidence of a dynamic predator-prey like relationship between circulating immune cell abundance and tumor size in patients that respond to immunotherapy. This relationship is not found either in patients that are non-responsive to immunotherapy or during chemotherapy. Single cell RNA-sequencing (scRNAseq) of serial peripheral blood samples from patients show that the strength of tumor-immune cell interactions is reflected in T-cells interferon activation and differentiation early in treatment. Thus, circulating immune cell dynamics reflect a tumor’s response to immunotherapy.

## Introduction

Immune checkpoint inhibitors can treat a wide range of cancers by targeting immune inhibitory pathways that cancer cells frequently coopt to avoid recognition and to regulate immune proliferation, survival, and effector functions (*1–11*). However, clinical response varies substantially, with approximately 40% of patients currently experiencing no objective benefit (*12, 13*). Numerous studies have investigated the role of tumor or tumor-associated immune cell phenotypes in response to immunotherapy (*14–19*). Patient responsiveness has been associated with increased tumor cell mutational load and antigen production (*20, 21*), and also with greater tumor-associated immune cell infiltration (*22*), signal production (*14*), and crosstalk (*23*). However, the consensus is that these markers are weakly associated with patient response (*24*). Furthermore, obtaining tumor tissue samples is challenging, especially if a tumor’s immunosuppressive phenotypes evolve over time.

Disease can regulate host immune cell abundance and signaling (*25–29*). Recently, it has been suggested that the frequency of specific peripheral blood immune cells can provide a non-invasive pre-treatment indicator of immunotherapy responsiveness, at least in melanoma cancer patients (*30*). As peripheral blood is easily accessible for serial analysis compared to tumor biopsies, a key question is whether circulating immune cells can serve as a surrogate measurement of a tumor’s interaction with the host immune cells and reflect response to therapy early in the course of treatment. If true, simple blood tests could be developed to guide patient specific clinical management decisions following the initiation of immunotherapy.

To address these questions, we have measured the strength of patients’ tumor-immune cell interactions, using a data driven ecological mathematical model of the concurrent dynamics of tumor and immune cell abundance. The strength of patients’ tumor-immune cell interactions was then related to immune cell phenotypes experimentally measured using single cell RNA-sequencing (scRNAseq). Fitting the tumor-immune cell interaction model to clinical tumor burden and immune abundance data revealed a consistently increased ability of responders’ immune cells to increase in abundance and indicated that improved tumor cells attack, drove decreased tumor burden. The increase in circulating immune cell abundance is concordant with a bolstered anti-tumor interferon signaling state of circulating immune cells and differentiation of T-cells to more cytotoxic states; as measured by scRNAseq. This combination of mathematical modeling and genomic analyses suggest that peripheral blood immune cell phenotypes reflect cancer-immune cell interactions and can reliably reveal patient responsiveness to immunotherapy.

## Results

### 1. Overview of trial and patient cohort

Patients with advanced GI cancers (colorectal, gastroesophageal, pancreatic and biliary) were enrolled in a single institution phase I trial (NCT02268825) of modified FOLFOX6 (mFOLOFX6) chemotherapy regimen followed by a combination of chemotherapy and anti-PD-1 immunotherapy (pembrolizumab) (**Fig. 1A**). Patient response was assessed according to the RECIST 1.1 guidelines, with responders showing complete/partial response (CR/PR) or stable disease (SD), and non-responders exhibiting progressive disease (PD) (Table S1-S2). Confirming our classification, 89% of responders survived more than 18 months after completion of treatment compared to only 26% of non-responders (**Fig. 1B**). As reported previously, the tumor’s PD-L1 expression was not strongly predictive of patient response (*24*). Single cell phenotypic insights (**Fig. 1C-D**) were linked to immune cell function by: i) mathematically modelling patients’ time courses of tumor burden and immune abundance, ii) fitting this model to the clinical data, iii) analyzing temporal changes in the growth rate of the tumor and immune cells and iv) relating patient specific model predictions to scRNAseq peripheral immune cell phenotype (**Fig. 1E**).

**Figure 1:**
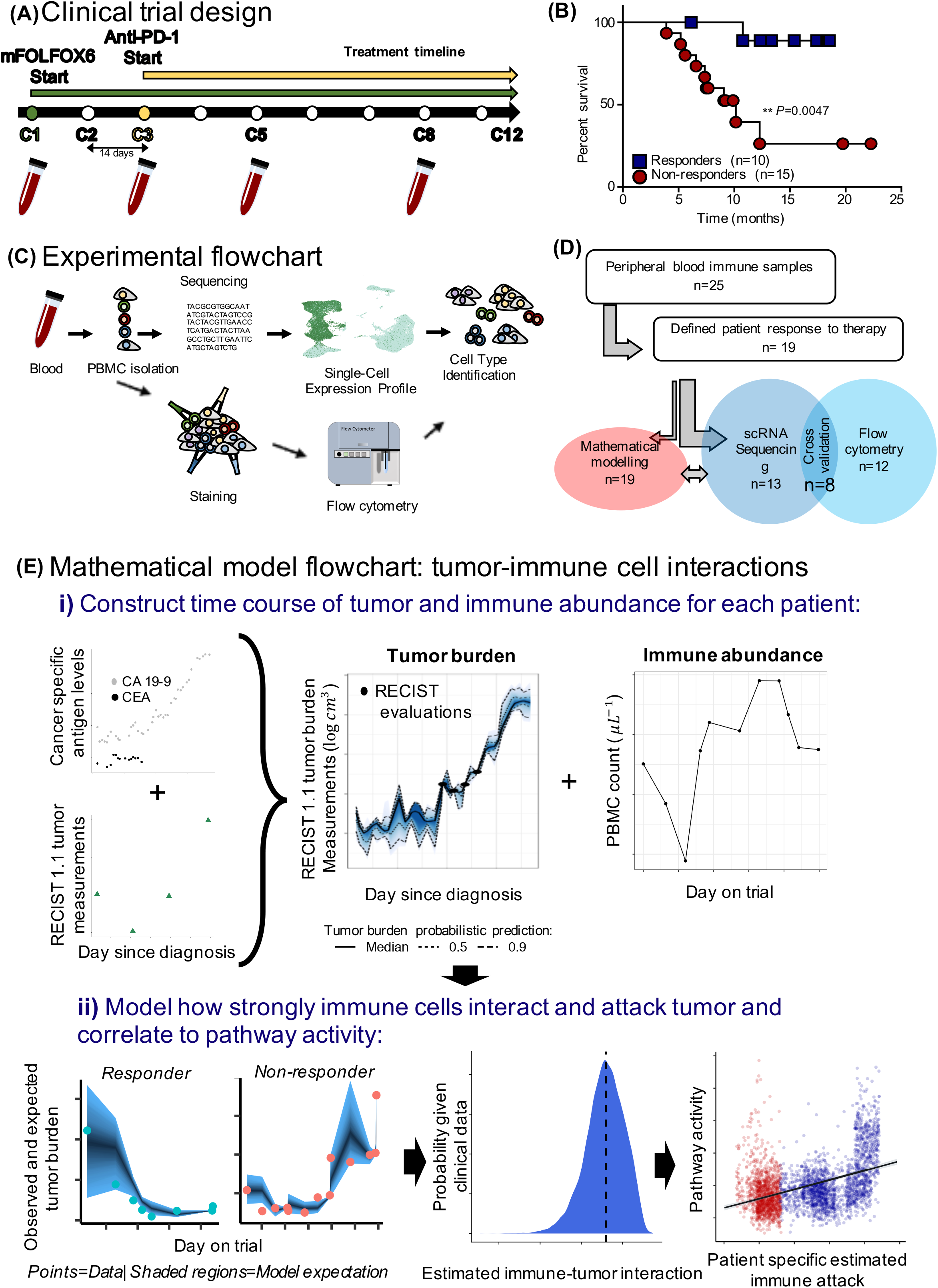
Overview of the clinical trial treatment strategy, patients’ classification, immune single cell analysis pipeline and tumor-immune interaction modelling. (**A**) Advanced gastrointestinal patients received mFOLFOX6 chemotherapy at the beginning of the trial for two 14-day cycles. From cycles 3 through 12, they received both mFOLFOX6 and anti-PD-1 immunotherapy. At baseline (cycle 1=C1), cycle 3 (C3), cycle 5 (C5) blood was collected and PBMCs were isolated and frozen. (**B**) Overall survival of responders and non-responders. (**C**) PBMC analyses using single-cell RNA sequencing and flow cytometry validation. (**D**) Flow chart of patient sample selection criteria, showing how patient samples were filtered and analyzed. (**E**) Mathematical modelling flow chart, depicting how i) clinical tumor burden data was synthesized and linked to concurrent measurements of PBMC abundance and ii) how a dynamic model of tumor-immune cell interactions, fitted to this data, allow inference of key biological processes (e.g. the ability of immune cells to kill tumor cells).

### 2. Patient specific immune function linked to immunotherapy success

Time courses of tumor burden and immune abundance (peripheral blood mononuclear cells: PBMC’s) were constructed for each patient (**Fig. 1Ei)**. Lymphocyte and monocyte abundance was strongly positively correlated with total immune abundance (Fig. S13), indicating a tight coupling of their population dynamics and motivating the modelling of total immune counts. Tumor burden was measured by combining information from cancer specific antigen biomarkers and RECIST 1.1 measurements of tumor size, using a Gaussian process latent variable model (SI Appendix). The changes in patients’ tumor burden and immune cell abundance during the trial were described mathematically by a dynamic model of cancer-immune cell interactions (**Fig. 1Eii**). In ecology, interactions between species, where the survival of one depends on attack by another, can be described using predator-prey equations. An adaptation of this ecological theory allowed us to describe the interactions between populations of tumor and immune cells within individual patients. We estimated the strength of this interaction, by statistically matching the changing frequency of immune cells and tumor size to model predictions. In the model, the tumor cells (T) are attacked by immune cells (I) and tumor cells induce increase immune cell recruitment. Chemotherapy (C) kills both tumor and immune cells, whilst PD-1i immunotherapy (P) impacts immune proliferation, recruitment and cytotoxic tumor activity (**Fig. 2A)**.

**Figure 2:**
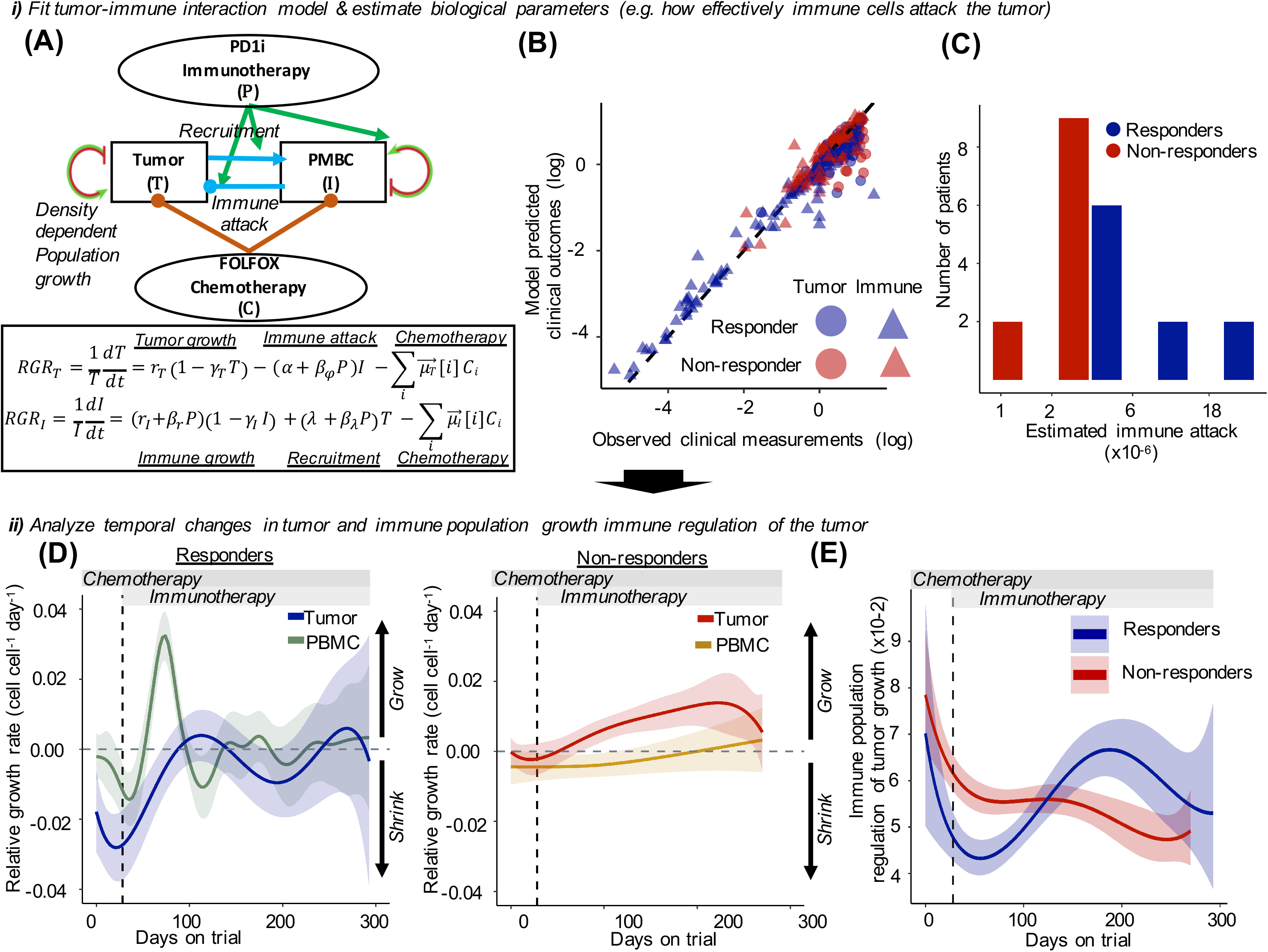
Patients’ immune cell function in attacking cancer cells and regulating tumor growth measured using a data driven tumor-immune cell interaction model. (**A**) Schematic of the mathematical model describing the strength of tumor-immune cell interactions and how their abundances change within a given patient over time. Blue arrows indicate recruitment (triangle tip) and attack interactions (circle tip) between cell types. Green arrows show how immunotherapy influences these interactions and immune population growth. Red arrows indicate chemotherapy effects. Curved arrows indicate intrinsic growth and density dependence within cell types. (**B**) Statistically fitting the model to clinical data allows an accurate description of observed tumor burden and PBMC abundance across patients and over time. (Dashed black line=1:1 model-data correspondence). (**C**) Histogram showing that responder patients consistently have immune cells with a higher ability to attack cancer cells. (**D**) Comparison of the speed of growth or decline of the tumor and immune cell populations during the trial, as measured by the relative growth rate of each component between observations. The distinct burst of immune activation in responders (LHS panel) and subsequent tumor decline was negligible in non-responders (RHS panel). Solid lines show mean trajectories and shaded regions signify model uncertainty intervals (vertical dashed line= start of immunotherapy, horizontal grey dashed line= stable population size). (**E**) Tumor-immune interaction model predictions of the ability of the immune cells of responders and non-responders to regulate the growth of the tumor during the trial.

Changes in tumor and immune cell abundance over time were accurately described by statistically fitting the mathematical model to the clinical data, using a Bayesian hierarchical approach (**Fig. 2B**). This analysis captured the biological differences between tumor and immune populations of responders and non-responders and the substantial variation between patients within these response categories. Key biological rates that were estimated included: a) how effectively immune cells attack the tumor and b) the impact of chemotherapy on tumor and immune populations. This identified the consistently improved ability of responder patients’ immune cells to attack the tumor, compared to non-responders (**Fig. 2C**).

The timing of most rapid growth/decline of tumor and immune populations were determined by analyzing the population’s relative growth rates (RGR= speed of population change, positive=growth, negative=decline) (**Fig. 2D-E**). The response dynamics were not dependent on the patient’s cancer tissue type. The tumor burden of the responders declined more rapidly during the chemotherapy phase and continued to decline (negative RGR) over time (**Fig. 2D)**. The exception is a time window around day 100 when the immune population was still increasing but the chemotherapy effect was generally decreased; once immune abundance reached a critical level, the tumor began to shrink once again and tumor burden remained substantially below the pre-treatment level for the duration of the trial. Interestingly, responders’ PBMC’s were also initially less abundant and more sensitive to chemotherapy (more negative RGR) (Fig. S14). However, their immune cell abundance was boosted following the addition of immunotherapy (**Fig. 2D**; spike in PBMC’s RGR around days 48-100). Their immune abundance then stabilized at this level or even increased gradually during the rest of the trial (overall positive RGR).

In contrast to responsive patients, the tumor burden non-responsive patients declined very little during the pre-immunotherapy chemotherapy phase, and only marginally in the first weeks of immunotherapy (**Fig. 2D**). Subsequently, tumor growth accelerated, and the tumor burden returned to the pre-treatment level within just 80-150 days. Further, non-responders exhibited a continual decline in immune cell number (negative RGR over most of the trial) and did not experience the immunotherapy induced boost in immune population growth following the addition of immunotherapy or benefit from immunotherapy. Model analysis showed that prior to immunotherapy, the responders’ immune populations less effectively regulated tumor growth (**Fig. 2E**). However, after immunotherapy induced the growth spike in the responders’ immune population, they became more effective at regulating tumor growth. In contrast, the ability of non-responders’ immune cells to regulate tumor growth declined continually during the trial and very little benefit of immunotherapy was detected.

### 3. Immune cell populations identified using scRNA-Seq profiles

To understand how phenotype changes of circulating immune cells related to the population dynamics and cell interactions (detailed above), we analyzed phenotypes of PBMCs isolated at 3 time points during the trial (**Fig. 1A, C**). Samples at cycle 1 (C1) provide the baseline before treatment, cycle 3 (C3) reflects treatment with only mFOLFOX6 chemotherapy, and cycle 5 (C5) reflect treatment with both chemotherapy and anti-PD-1 immunotherapy. A total of 13 patients (responder n=7, non-responder n=6) were analyzed by scRNAseq (**Fig. 1C**). The transcriptional profile of 70,781 immune cells was obtained, revealing a diverse set of 35 cell types. All major PBMC lineages were identified using canonical gene expression markers and analysis of a uniform manifold approximation and projection (UMAP) (**Fig.3**, Fig. S1-S3, Table S3).

The cell type annotations were validated by comparing our transcriptional profiles and corresponding annotations with published studies of PBMC’s (*31*) and tumor infiltrating immune cells (*32*). We found that 96.5% of T-cells from the PBMC database and 94.1% of T-cells from the tumor infiltrating dataset were correctly predicted using a machine learning classifier trained using our annotations (**Fig. 3B**, Fig. S3). A similarly high agreement was found between our annotations and published annotations when examining cell type specific marker genes and comparing the cell type connections (**Fig 3B**, Fig. S3-S4). As a final validation, we profiled 8 patients (6 responders, 2 non-responders) with both scRNAseq and flow cytometry (Fig. S5). An approximate 1:1 correspondence was found between the abundance of immune cell types obtained using each method (Fig. S6). Immune cell numbers were quantified in two ways: i) the frequency of cells refers to the percentage of cells in a sample, ii) the abundance refers to the measured number of cells per unit of peripheral blood.

**Figure 3:**
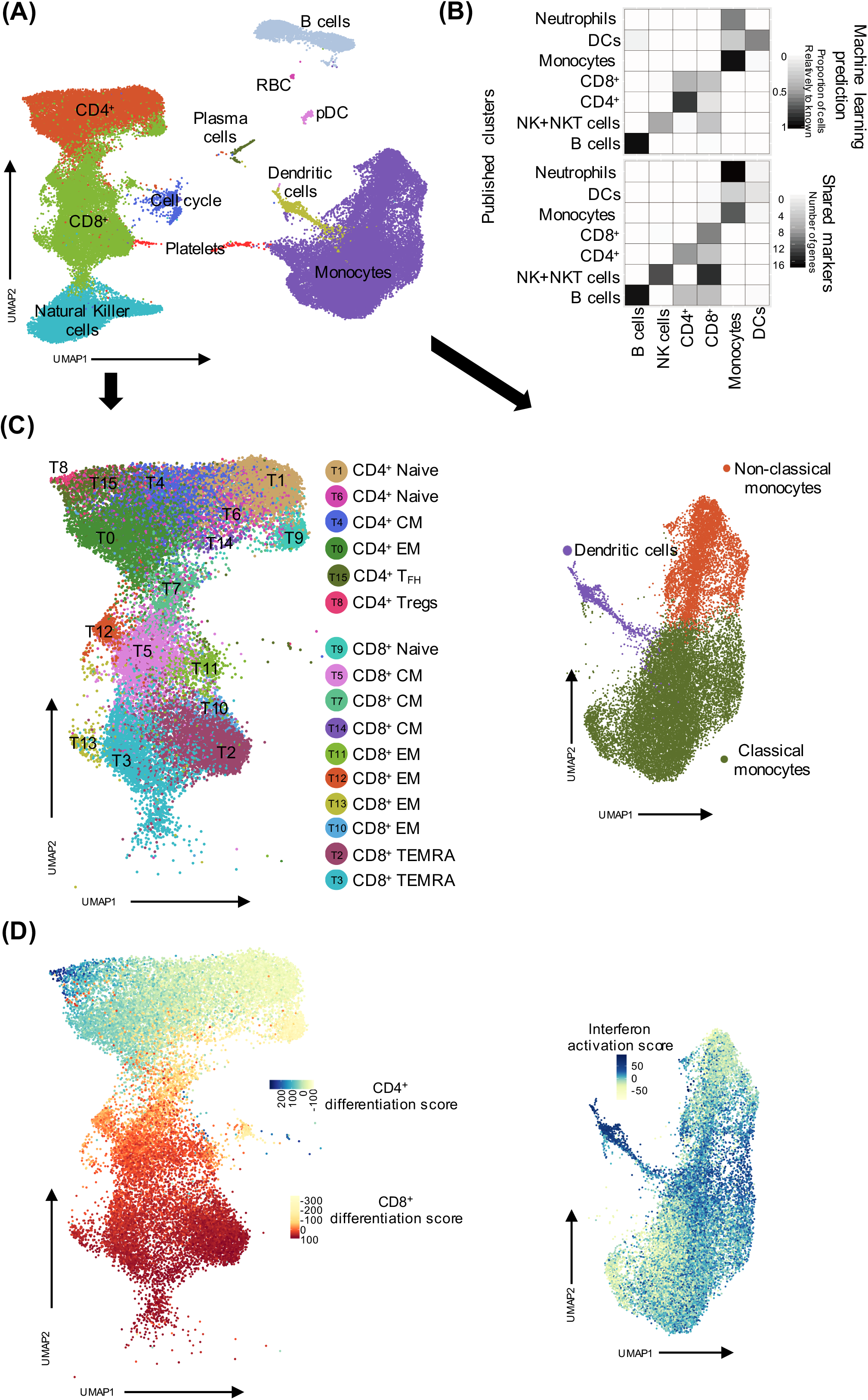
Validated classification of immune cell types, T-cells and monocyte subtypes and identification of the major phenotypic variation within these populations. (**A**) Uniform Manifold Approximation and Projection (UMAP) of the single cell RNA sequencing (scRNAseq) data of all patient’s PBMC’s across analyzed time points. Major PBMC types are labeled (RBC= red blood cells, pDC= plasmacytoid dendritic cells). (**B**) The agreement between our predicted clusters and public classifications of cell types annotated in two published datasets. Top panel (machine learning prediction): the distribution of immune cells in public datasets predicted to our annotation clusters by Random Forest learner using our predicted clusters as a training set. Bottom panel (Shared marker genes): the number of shared genes between public datasets and our predicted clusters (SI Appendix; NKT=Natural killer T-cells, DCs=Dendritic cells). (**C**) UMAP identification of CD4^+^ and CD8^+^ T-cell subclusters (T_FH_ = Follicular helper) and monocyte subtypes. (**D**) UMAP representing phenotypic gradients of CD4^+^ differentiation (top of left subplot: lowest score at right and highest to the left), CD8^+^ cytotoxic differentiation (bottom of left subplot: lowest score towards the top right and highest at the bottom) and monocyte interferon activation.

### 4. Signaling activation in responders’ T-cells upon initiation of immunotherapy

Signaling dynamics upon initiation of immunotherapy were examined through single cell pathway activity analysis, using single sample Gene Set Enrichment Analysis (ssGSEA) scores (33) of C2-level and Hallmark pathway signatures (*34, 35*). Pathway differences before therapy, during chemotherapy and during the early-immunotherapy phase of the trial were identified using a random effects linear modeling framework (**Fig. 4**). This approach partitioned the effects of chemotherapy and immunotherapy on pathway activity while accounting for individual variability in expression. The statistical significance of P-values was corrected using Holm’s conservative multiple comparison correction procedure.

**Figure 4:**
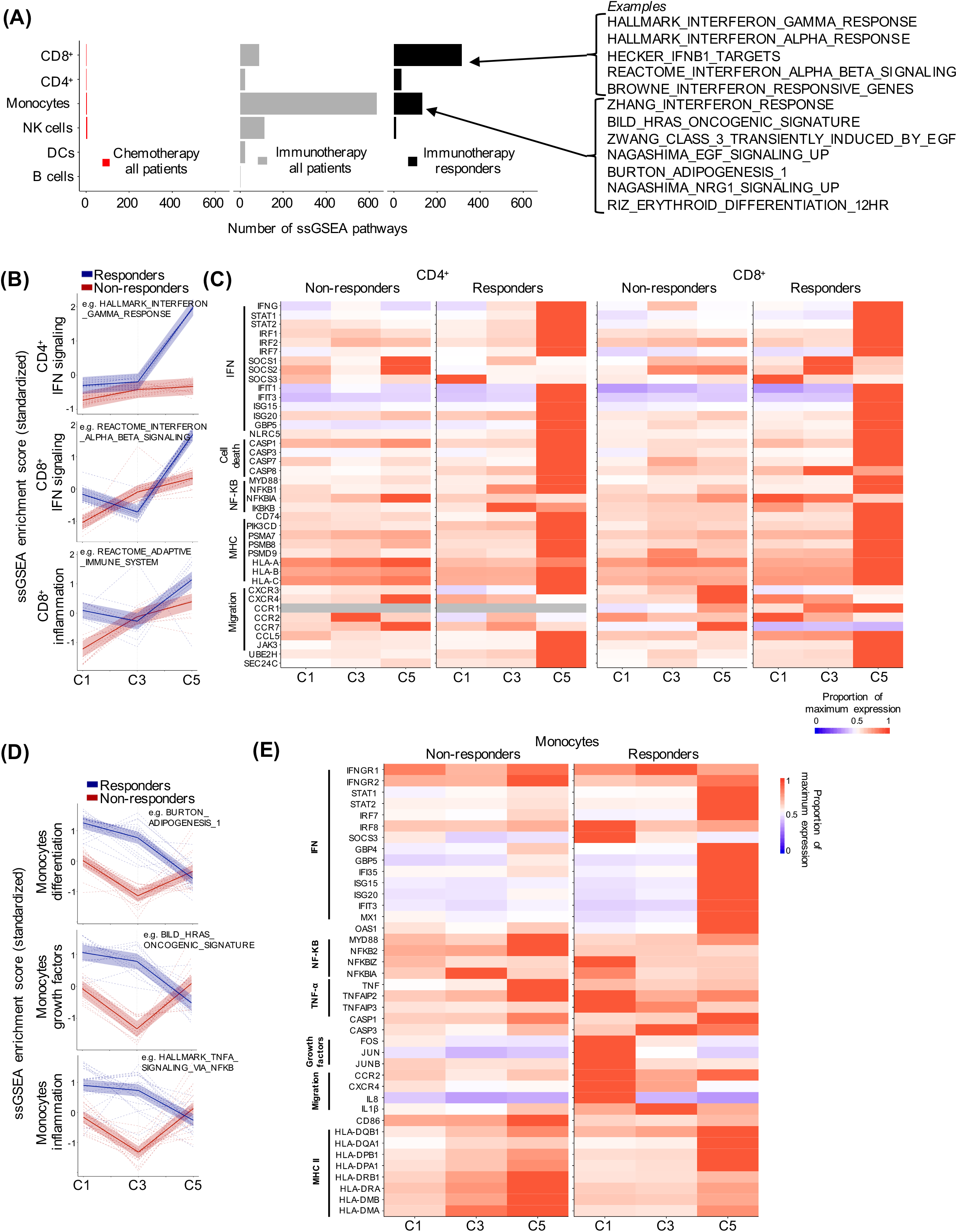
Pathway signaling activation of multiple immune cell types in responders but not non-responders following initiation of immunotherapy. (**A**) The number of molecular pathways impacted by chemotherapy and PD-1 immunotherapy and whether PD-1 immunotherapy effects are specific to responders (black bars) or common across patients. The “chemotherapy all patients” panel shows the numbers pathways changing expression between time C1 and C3 in different-cell types. The “immunotherapy all patients” panel shows the numbers of pathways showing trends in expression between C3 and C5 which are common to responders and non-responders. Finally, the “immunotherapy responders” panel shows the numbers of pathways with trends in expression that are unique to responder patients. Pathways with very differing trends in responders and non-responders are exemplified on the right side. (NK = Natural killer, ssGSEA=single sample Gene Set Enrichment Analysis). (**B**) Interferon and inflammatory signaling of CD4^+^ and CD8^+^ T-cells is upregulated in responders more than non-responders. GSEA pathway categories reflect the most enriched types of pathways for each cell type. Individual GSEA pathways exhibiting differential trends in expression between responders and non-responders are shown (dashed lines). Overall trends of pathways within each cellular process (solid lines) and variation (shaded regions) are overlaid (IFN=Interferon). (**C**) Heatmap of changes in gene expression of responder and non-responder CD4^+^ and CD8^+^ T-cells over time. IFN, cell death, NF-κB, MHC (major histocompatibility complex) I & II and migration signature genes are displayed as the proportion of maximum level of each gene. Genes not detected in a cell type are shaded grey. (**D**) Differences in inflammatory signaling, differentiation and growth factor production between the monocytes of responders and non-responders showing overall trends of pathways within each cellular process (solid lines) and variation (shaded regions). Trends of pathways exhibiting differential expression patterns in responders and non-responders are indicated by dashed lines. (**E**) Heatmap of changes in gene expression of responder and non-responder monocytes over time. Interferon, cell death, NF-κB, TNF-α, growth factors production, and migration signature genes are displayed as the proportion of maximum level of each gene. Statistical significance of differences between responders and non-responders was determined for each gene and corrected for multiple comparisons. C1= cycle 1: baseline, C3= cycle 3: chemotherapy mFOLOFX6 regimen, C5= cycle 5: Chemotherapy + anti PD-1 immunotherapy. One cycle = 14 days.

Overall, immune cell gene expression was not greatly altered during chemotherapy treatment (**Fig. 4A**, left panel). In contrast, after the start of anti-PD-1 treatment, there were a subset of pathway changes common to both responder and non-responder’s monocytes and T-cells (**Fig. 4A**, middle panel). Further, a majority of signaling changes were identified that were specific to responders (**Fig. 4A**, right panel, Table S3). For each immune cell type, the most significantly altered GSEA pathways were classified into categories reflecting major biological processes.

Strikingly, interferon signaling pathway activity was significantly upregulated in CD4^+^ and CD8^+^ T-cells of responder patients following the initiation of anti-PD-1 treatment (C3-C5) (CD4^+^: *t*=19.00, *p*<0.001, CD8^+^: *t*=16.00, *p*<0.001) **(Fig. 4B**, Fig. S7). CD8^+^ T-cells of non-responders showed a lesser upregulation of interferon signaling after the start of anti-PD-1 (t=7.61, p<0.001), while CD4^+^ T-cells show no such increase. Upon initiation of immunotherapy, a range of interferon related genes were upregulated in the CD8^+^ and CD4^+^ T-cells of just the responders (**Fig. 4C**, Fig. S8). Responders’ CD8^+^ cells showed greater upregulation of the IFN-γ gene (p<0.01) and IFN target genes (IRF1/2/7, STAT1/2 and interferon-stimulated genes (Table S4). In contrast, non-responders’ CD4^+^ and CD8^+^ T-cells had greater upregulation of IFN repressing genes (e.g. SOCS1 and SOCS2) (p<0.05), indicating impaired transduction of IFN signaling upon anti-PD-1 treatment (*36*). Inflammatory response pathways were also upregulated in T-cells of responders (**Fig. 4B**), including CD8^+^ T-cells of responders prior to the onset of any treatment (*t*=5.14, *p*<0.001) and after addition of anti-PD-1 (*t*=3.8, *p*<0.001). Inflammatory genes induced with anti-PD-1 include major histocompatibility complex (MHC class I/II) sorting and processing genes (e.g. CD74, HLA-A/B/C and PSM) as well as NF-κB pathway genes (NFKB1, IKBKB, MYD88) in responders’ CD8^+^ and CD4^+^ T-cells (**Fig. 4C**, Table S4). The NF-κB activation of responders’ T-cells may suggest a shift to a pro-survival state. Overall, this shows the activation of these peripheral cells and the increased signal transduction in responders.

### 5. Patients responsive to therapy exhibit changes in monocyte signaling during treatment

Monocytes also exhibited different phenotypes in responders versus non-responders but with distinct signaling changes from those of T-cells. Before treatment (C1), monocytes from responders had significantly higher activation of three pathways representing related but distinct measures of monocyte developmental states: growth factor production (*t*=9.2, *p*<0.001), inflammation (*t*=6.1, *p*<0.001), and differentiation (*t*=6.3, *p*<0.001) (**Fig. 4D**). While chemotherapy decreased each of these pathway scores in both responders and non-responders, patients responsive to anti-PD1 treatment exhibited a significant reduction in all three pathways after anti-PD-1 treatment (*p*<0.001 for each pathway) while non-responders showed a significant increase (*p*<0.001 for each pathway). During immunotherapy, responders and non-responders’ monocytes showed specific gene dysregulation of: growth factor, IFN, TNF, NF-κB, and MHC genes (**Fig. 4E**, Fig. S9). In addition, genes promoting the migration and recruitment of other immune cells types were initially upregulated in responders’ monocytes (CXCR4, CCR and CCL family members) (*37*) (Table S4). Overall, monocytes showed pretreatment differences in signaling and divergent developmental trajectories in responders versus non-responders. Activation of monocytes after the start of anti-PD-1 may reflect responses to the upregulation of IFN and cytokine gene expression observed in responders’ T-cells.

### 6. During therapy, T-cells of responders differentiate, while non-responder CD8 T-cells lose cytotoxicity

The major phenotypic differences within each immune type were identified, using pseudotime reconstruction of scRNAseq profiles (Fig. S10). By overlaying the cellular phenotype scores onto a UMAP of the expression profile, we validated that the phenotypes reflect the key sources of transcriptional variation within immune cell types (**Fig. 3D**). The CD4^+^ T-cell phenotypic gradient captured the continuum of differentiation from naïve to effector helper T-cells (**Fig. 3D** left panel). Similarly, the CD8^+^ T-cell phenotype gradient captured differentiation from a naive to highly cytotoxic state. In both cases, naive, central memory, and effector T-cell subtypes aligned clearly along the continuous phenotype gradient and in the expected order.

We next evaluated the distribution of T-cell phenotypes in the peripheral blood of responders and non-responders and examined how they shifted during the course of therapy (**Fig. 5A-B)**. Before treatment (C1), responders had a higher frequency of undifferentiated (naive) CD4^+^ T-cells, which may have been symptomatic of the tumor-mediated immune suppression (**Fig. 5B**). In contrast, non-responders had more differentiated CD4^+^ T-cells, especially CD4^+^ EM cells (*t*=−7.5, *p*<0.001) (**Fig. 5B**). This difference remained following the onset of chemotherapy (C3); however, after immunotherapy (C5), the CD4^+^ T-cells of responders showed a significant shift towards increased differentiation (*t*=9.9, *p*<0.001) and converged with non-responders (Fig. S11a). Interestingly, responders had a higher frequency of cytotoxic differentiated CD8^+^ T-cells than non-responders, both before and during treatment (**Fig. 5B**, Fig. S11b) (*F*=16.8, *p*<0.001). With the addition of anti-PD-1, responders’ CD8^+^ T-cells became even more cytotoxic (*t*=3.9, *p*<0.001), while non-responder’s CD8^+^ T-cells shifted to a less cytotoxic state (*t*=−4.0, *p*<0.001).

**Figure 5:**
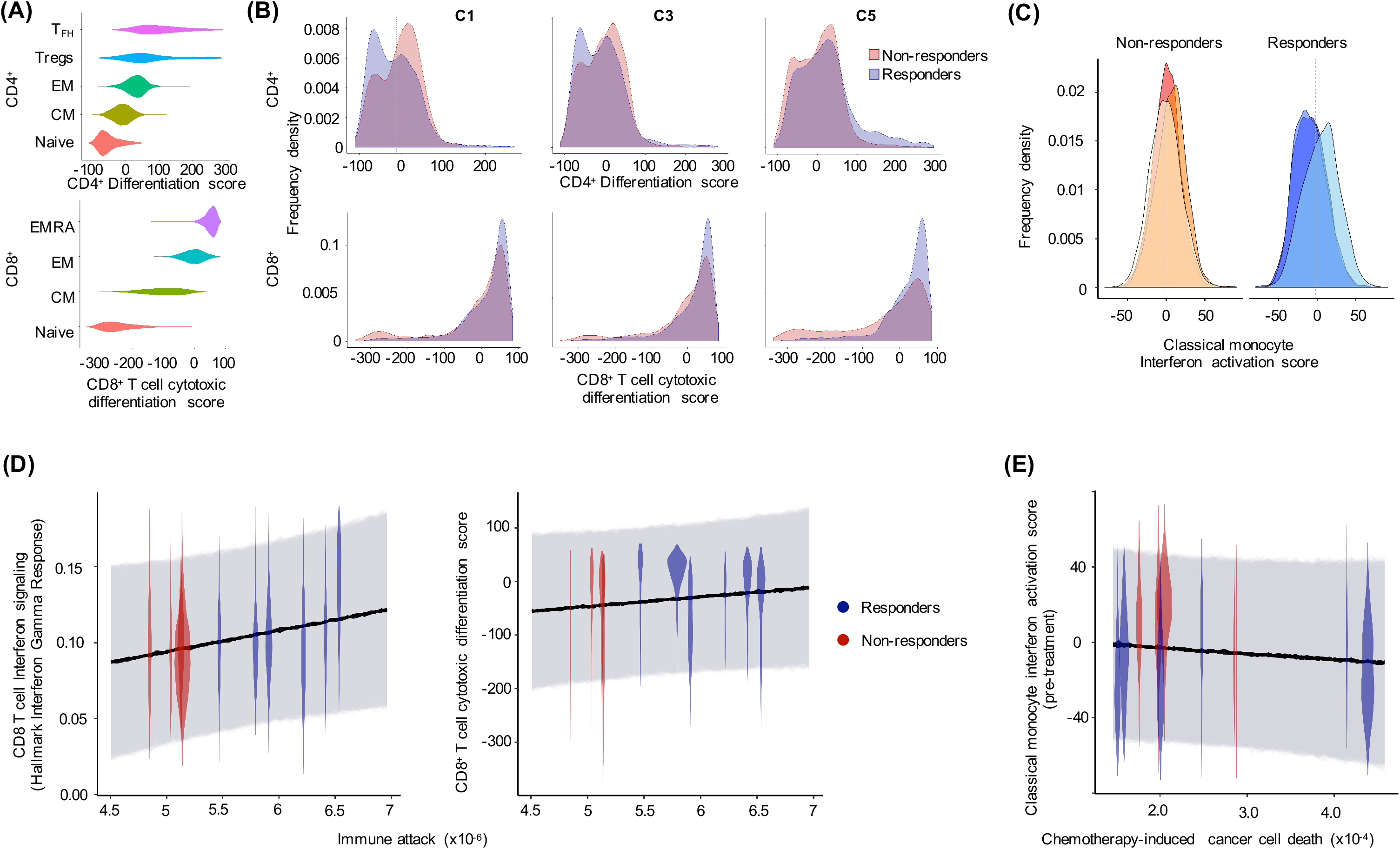
Peripheral blood immune cell phenotypes linked to patients’ immune cell function and immunotherapy responsiveness. Responsiveness to immunotherapy depends on circulating memory T-cell differentiation and monocyte interferon activation prior to therapy. (**A**) Comparison of CD4^+^ and CD8^+^ T-cell subtype differentiation scores (all subtypes differ with a Tukey test) (EM=Effector memory, EMRA=Effector memory CD45RA^+^, CM=Central memory). (**B**) Frequency of CD4^+^ and CD8^+^ T-cells with different states of differentiation/cytotoxicity in responders and non-responders at each treatment time point. (**C**) Frequency of monocytes with different interferon (IFN) activation states in responders and non-responders at each time point. (**D**) The ability of patients’ immune cells to attack cancer cells and also the tumor’s sensitivity to chemotherapy was linked to immune cell signaling and differentiation phenotypes. For each patient, the single cell variability in immune cell phenotypes are presented as individual violins densities. Black line indicates the relationship between a patient’s average immune cell phenotype and the strength of immune cell attack /chemotherapy sensitivity. Shaded regions = credible intervals for the predicted range of phenotypes of 95% of the immune cells, given the strength of immune cell attack/chemotherapy sensitivity.

### 7. Monocytes of responders were activated after the start of anti-PD-1 therapy and the frequency of classical monocytes was associated with response

Within monocytes, the expression of interferon response genes was the major axis of phenotypic variation (**Fig. 3C-D** right panel). Monocytes with high interferon response scores (including dendritic cells) had upregulation of IFN stimulation genes (e.g. IFIT1/3, PSME2, and ISG15) and higher MHC class II expression (e.g. *HLA.DPA1, HLA.DPB1,* and *HLA.DMA*). In contrast, cells with low interferon scores had upregulation of proliferation (e.g. FOS, JUN, and JUNB), differentiation (e.g. BTG1, RGS2, and DDX17), inflammation (e.g. SELL, S100A12, and CD36) and migration (e.g. VCAN and VIM) genes. After immunotherapy, monocytes with the highest interferon score became prevalent in responders (*t*=15.463, *p*<0.001) (**Fig. 5C**, Fig. S14d). Responder patients shifted from having the lowest to the highest average level of interferon activation and MHC class II gene expression (Fig. S12). In contrast, the distribution of interferon response in non-responder monocytes remained relatively constant across the trial period.

### 8. Linking immune function and phenotypes of the peripheral blood

Finally, we linked the patient specific estimates of immune attack and chemotherapy sensitivity to the single cell transcriptomic observations of increased immune cell signaling and phenotypic differentiation states in responders (**Fig. 5D-E**). Patients whose immune population had a greater ability to attack tumor cells and response to immunotherapy were found to have CD8^+^ T-cells with higher activity of interferon gamma signaling pathways and more differentiated cytotoxic CD8^+^ T immune cells (**Fig. 5D**). Finally, patients whose monocytes showed lower activity of interferon gamma response pathways (classical monocyte differentiation score) before treatment had tumor cells that were significantly less sensitive to chemotherapy (**Fig. 5E**).

## Discussion

Our findings indicate that peripheral blood immune cell phenotypes reflect the strength of tumor-immune interactions before or early in the course of immunotherapy, and these phenotypes are indicative of patient responsiveness. By combining scRNAseq analysis of peripheral immune phenotypes with dynamical models of patient specific clinical data, we linked peripheral immune cell phenotypes with the strength of patients’ tumor-immune cell interactions. Increased interferon signaling and differentiation of T-cells was related to an increased ability of immune cells to attack cancer cells, regulate tumor growth and drive patient responsiveness to anti-PD-1 therapy. These results provide motivation for studies interrogating the utility of peripheral blood phenotypes as a biomarker of patient responsiveness to therapy.

Although mathematical modeling has provided important insights into cancer-immune cell interactions and cancer immunotherapy, models incorporating patient specific clinical or phenotypic data had not previously been developed (*38–45*). Previous theoretical models that do not include patient data have described the potential for cancer-immune interactions to act as “predator-prey like” systems (reviewed in (*45*)). This study is a step forward in that it uses temporal clinical and single cell immune phenotyping for data driven ecological modelling of patient-specific responses during treatment.

The cancer-immune interaction model predicts that in general, patients whose tumors have an immunosuppressive phenotype (e.g. expressing high levels of PD-L1) will have a lower immune cell count prior to treatment, as immune activation and proliferation is inhibited. Hence, we expect that patients with a low PBMC abundance should benefit most from anti-PD-1 immune re-activation therapy. In agreement, we observed significantly lower PBMC abundances in responders at the onset of therapy (Fig.S15a). These patients showed gradually increasing immune counts during therapy, in contrast to declines observed in non-responders. Model analysis indicated that, at the onset of the trial, the immune cells of responders had a substantially weaker effect of tumor regulation compared to those in non-responders, primarily due to the low immune cell count (Fig.S15b). During immunotherapy, the responders’ immune population gradually increased and their tumor regulatory effect increased towards the level of the non-responders. This leads to the prediction that, unlike chemotherapy, the tumor’s response to immunotherapy will be delayed. This is a general prediction that emerges from predator-prey models. Due to fewer immune cells present and few cancer antigens being presented to initiate further immune response prior to therapy, several rounds of the cancer-immune response cycle are needed for the immune population to rebuild following PD-L1 suppression.

Our model also predicts that chemotherapy acts as a double edged sword when used as a combination therapy with immunotherapy. It has the positive effect of inducing tumor cells death and promoting immune cell recruitment; however, it also kills immune cell progenitors, reducing the active immune cell abundance. Therefore, too high a chemotherapy dosage may inhibit the effectiveness of immunotherapy, whilst too low a level may not promote immune re-activation.

The analyses of T-cell and monocyte signaling states, before and during therapy, suggest that circulating immune cells rapidly shift phenotypes during the treatment in GI cancer patients. We suggest that this peripheral immune signaling activation is a valuable early marker of patient responsiveness. The interferon surge after initiation of anti-PD-1 therapy, seen only in responders’ T-cells and monocytes, indicates that treatment with anti-PD-1 is promoting differentiation and activation of T-cells, resulting in antitumor activity, cytokine release, and stimulation of the immune system. In particular, only responders CD8^+^ T-cells upregulate IFN-γ signaling and immune cell activation and anti-tumor effect (*46*). Despite PD-1 blockade, non-responders’ immune cells were not fully activated, indicating that they struggle to detect cancer cells. Possibly, low cancer antigen release, reduced activation of antigen presenting cells and T-cells, and prevented initiation of an immune response. Additional studies support an interaction of chemotherapy with immunotherapy in some settings (*47–51*). Using our scRNAseq time courses, we also detected that immunotherapy induces a shift to a more differentiated CD4^+^ T-cell state. Long term chemotherapy may increase the production of PD-1 expressing regulatory CD4^+^ EM cells, diminishing pembrolizumab availability to tumor-specific CD8^+^ T-cells (Fig.S16).

Additionally, patients may have been non-responsive because cancer cells had PD-1 independent resistance mechanisms of immune avoidance. Indeed, we found that non-responder’ classical monocytes had low MHC II receptor expression suggesting lower antigen recognition and presentation. They also developed a more immunosuppressed phenotype, with upregulation of CD86, a ligand of both PD-1 and CTLA-4, and CD28, a costimulatory signal for activation of T-cells. Contrastingly, under anti-PD-1 therapy responders’ monocytes showed activation of costimulatory immune function (upregulated ISG and MHC).

Overall, we find that the abundance, signaling activity and differentiation state of peripheral immune cells reflect tumor-immune cell interactions and patient response to immunotherapy. The combination of total PBMC abundance and the relative infrequency of differentiated/ activated effector T-cells likely provides a non-invasive upfront marker of therapeutic responsiveness. Models of tumor-immune cell interactions, which use clinical and phenotype data, allow quantification of the immune system’s effectiveness in regulating tumor growth and demonstrate the potential of using peripheral blood-based models to assess the dynamics of the immune and tumor cell interactions during treatment.

## Materials and Methods

### Study design

Cryopreserved peripheral blood mononuclear cell (PBMC) samples from patients with advanced (stage 3/4) gastrointestinal cancers were collected from patients in a clinical trial (NCT02268825), and were treated with modified FOLFOX6 regimen every 2 weeks (i.e. 1 cycle) until disease progression, death, or completion of the study. After 4 weeks of mFOLFOX6 (cycle 3), pembrolizumab (200 mg IV every two weeks) was added to mFOLFOX6. Before treatment and then every two weeks, patients’ blood was collected and PBMCs were isolated and cryopreserved. All human biological samples were collected after written informed patient consent and ethics committee approval, following federal and institutional guidelines. The University of Utah Institutional Review Board and the Huntsman Cancer Institute Protocol Review and Data and Safety Monitoring Committee approved and monitored this study.

The primary outcomes of this phase I study was safety and dose limiting toxicities. Patients were excluded if they had active infection, autoimmune disease, or were on chronic systemic steroids or immunosuppressant’s. Samples from 13 patients (responder n=7, non-responder n=6) were used for scRNAseq analysis at C1, C3 and C5 time points. Samples from eight patients were utilized for both FACS and scRNAseq analysis (responder n=6, non-responder n=2), to validate the consistency of inferences. Single cell transcriptional profiling provided information for a total of 70,781 cells from 13 patients.

Clinical response was measured by computed tomography scans and assessed according to RECIST1.1 and immune-related response criteria (irRC) every 12 weeks. Responders were defined as patients with clinical benefit at 24 weeks (complete response (CR), partial response (PR) or stable disease (SD)). Non-responders included patients with progressive disease (PD defined as > 20% increase in tumor volume or appearance of new metastatic lesions) between 12 and 24 weeks after the trial began. Median of previous history of chemotherapy treatment for responders was 101 days and 42 days for non-responders (**Table S1**).

### Single-cell RNA sequencing and annotation

PBMC samples analyzed using a Chromium 10X Cell Instrument (10X Genomics) (1200-2000 cells/sample) and sequenced on an Illumina HiSeq 2500 with 2×125 paired-end reads. Raw BCL sequencing files were processed using Cell Ranger Single Cell Software Suite and samples were aligned to hg19 using the STAR aligner (*52*). Count tables were generated for 70,781 cells and used as input into Seurat v2 (*53*). No batch effects were found corresponding to time, patient or cancer (Fig.S2 b-d).

To identify cell types, variable genes (n=1000) and non-overlapping known immune cell marker genes (n=1480) were used for PCA (*54–56*). The first 25 PCs captured significant variation, based on Seurat’s jackstraw analysis, and were used for graph-based clustering and UMAP visualization (*57*). Major T-cell clusters were identified using *CD3D*, *CD4* and *CD8* expression along with 500 T-cell specific variable genes and 273 known T-cell markers (*56*). Differential expression markers for each cluster were generated using MAST(*58*). Pathway ssGSEA enrichment scores were generated using the R package GSVA 1.30.0 (*33*). Immune cell annotations were verified using two public datasets (*31, 32*) (SI Appendix, Fig. S3-4) using training and classification to measure similarity of annotation.

### Identifying gene set expression differences between responders and non-responders

Differences in the gene set expression of immune cell types were examined between responder and non-responder patients (R). For each immune cell type, we examine the changes in pathway (X) expression over time (T) and with the addition of the anti-PD-1 (P). A random effects model with the following linear predictor (η) and error structure was constructed for each pathway:

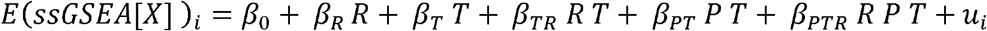

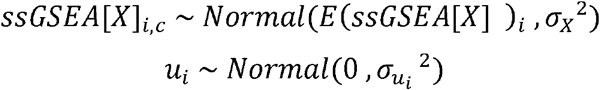

Initial differences in gene set expression between immune cells from responders and non-responders, at the pre-treatment time point (C1), were captured by the group-specific intercepts (*β_0_ vs. β_R_*). Differential trends in expression over the first 5 treatment cycles were described by the group specific slope terms of responders and non-responders (*β_T_ vs. β_TR_*). Differential effects of the addition of anti-PD-1 on gene expression, over cycle C3-C5, were described by the group specific anti-PD-1 treatment effect terms of responders and non-responders (*β_PT_ vs. β_PTR_*).

Background individual variability in gene expression, independent of therapy impacts, were accounted for by allowing the model intercept to vary among patients (*u*_*i*_). Significant differences in: A) initial pathway scores, B) temporal trend and C) anti-PD-1 treatment effects between non-responders and responders were assessed using likelihood ratio tests. Multiple comparison corrections were made using Holm’s p-value correction.

### Quantifying immune cell phenotypes

Major axes of phenotypic variation were identified separately for CD4^+/^CD8^+^ T-cells and monocytes using affinity-based pseudotime reconstruction of cell states (*60, 61*). This allowed the description of continuous spectrums of cellular states, as is produced by differentiation and activation processes (SI Appendix). These phenotypic axes were validated using comparisons to PCA, zinbwave and UMAP dimension reduction (*57, 63*). Random effects linear regression was used to test the statistical differences in immune population phenotype distributions between responders and non-responders, whilst accounting for patient-specific random effects.

### Modeling and measuring tumor-immune cell interactions

#### Overall measures of tumor burden

We assessed the strength of tumor-immune cell interactions and the predictability of responsive to therapy by fitting a coupled tumor-immune population model to clinical patient data (SI Appendix Dataset S1). For each patient, a time series of tumor burden was first constructed, by combining RECIST 1.1 measurements, from CT scans, with information from tumor burden biomarkers (CA 19-9 and CEA), using a Gaussian process model (*64*). Gaussian process models probabilistically combine these tumor burden data sources, allowing inference of tumor burden (SI Appendix).

#### Tumor-immune interaction model

The dynamics of tumor and immune cell abundance were coupled with the immunotherapy and chemotherapy dosing schedules, using a patient specific tumor-immune population dynamic model. The ecologically inspired model (Equ.1) describes the patient specific changes in tumor (T) and immune cell (I) abundance over time. Over short periods of time, the increase or decrease in tumor and immune cell abundance was measured by the populations relative growth rate (*Rc R*_*T*_ for tumor and *Rc R*_*I*_ for immune cells). Positive RGR values indicate population growth, whilst negative values show population decline. The data driven model decomposed this population growth rate into effects of different concurrent biological processes. Tumor and immune cells interact in two main ways, with tumor cells being attacked by immune cells (*α*) and also inducing increased immune cell recruitment (*λ*). Therapeutic dosing impacts the cell populations and the strength of their interactions, with chemotherapy (C) killing both tumor 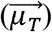 and immune cells 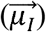, whilst PD-1i immunotherapy (P) influences immune proliferation (*β*_*r*_), recruitment (*β*_*λ*_) and cytotoxic tumor killing activity (*β*_*φ*_). Both tumor and immune cells experience density dependent population growth (*γ*_*T*_ & *γ*_*I*_), reflecting competition for resources or growth stimulating molecules. This leads to the equations:

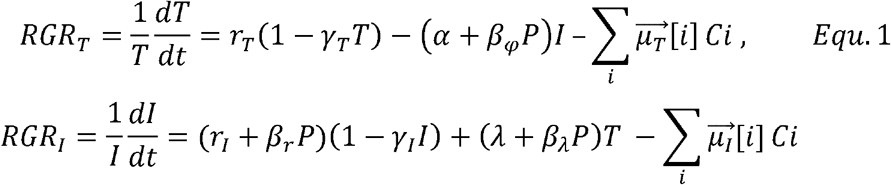

We simultaneously fitted this model to all of the patients’ time course tumor and immune data, and accounted for the differing dosages and timings of therapy. To capture inter-patient biological differences, patient specific parameters were assumed to be drawn from a hyper-distribution of parameters, creating a hierarchical model structure. Model parameters were estimated using Bayesian inference in Stan (*65*).

### Linking immune phenotypes and model estimated biological processes

Immune cell phenotypes were related to the model estimates of: a) the effectiveness of immune cells at attacking tumor cells and b) the tumor cell sensitivity to chemotherapy. These biological estimates of immune and chemotherapy function (X) were regressed against the peripheral immune cell phenotypes identified in: i) the GSEA pathway analysis and ii) the pseudotime analysis of the major phenotypic variation within cell types. For each phenotype, the significance of the relationship between single cell peripheral immune phenotypes (Y) and immune/chemotherapy function (X) was assessed. A patient specific intercept was added to account for non-independence of cell phenotypes within a patient. The random effects regression model was simply:

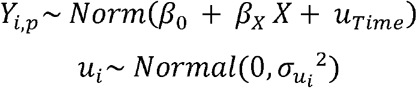

The significance of the relationship between peripheral phenotypes and immune/chemotherapy function was assessed using a likelihood ratio test, with the sample size corrected for the non-independence of data points.

## Acknowledgments

We thank the anonymous patients used in our study. We acknowledge S. Brady and B. Nelson for helpful feedback. The High-Throughput Genomics Shared Resource and was supported by the NIH Award Number P30CA042014. The content is solely the authors responsibility and does not necessarily represent the official views of the NIH. J.T.C. was supported by a Cancer Prevention Research Institute of Texas Core Facility Support Award (RP170668). A.H.B, J.T.C, J.G., P.C., P.W., X.L, P.J.M., J.A.M., F.A. were supported by the National Cancer Institute of the National Institutes of Health under award number U54CA209978.

## Author contributions

J.G., P.W., L.P. contributed to study design, coordination, wrote the manuscript, processed and analyzed the data. X.L., P.C., D.S., P.J.M., J.T.C., F.A., J.A.M. contributed to manuscript writing. N.L., X.L. and J.T.C contributed to data processing and analysis. J.A.M., G.Sh., P.J.M. coordinated sample collection and single-cell isolation. M.R. and G.Su. performed flow cytometry and contributed to the analysis. D.S, S.S supervised the clinical trial and compiled patient clinical data. A.H.B. conceived the study, contributed to study design, coordination, manuscript writing and data analysis.

## Data and materials availability

Raw single cell RNA-seq data have been deposited in GEO under accession code GSE130157. Tumor and immune abundance clinical time courses are provided in Dataset S1.

## References

1. M. Loos, et al., Clinical significance and regulation of the costimulatory molecule B7-H1 in pancreatic cancer. Cancer Lett 268, 98–109 (2008).

2. T. Nomi et al., Clinical significance and therapeutic potential of the programmed death-1 ligand/programmed death-1 pathway in human pancreatic cancer. Clin Cancer Res 13, 2151–2157 (2007).

3. Y. Ohigashi et al., Clinical significance of programmed death-1 ligand-1 and programmed death-1 ligand-2 expression in human esophageal cancer. Clin Cancer Res 11, 2947–2953 (2005).

4. E. Oki et al., Protein Expression of Programmed Death 1 Ligand 1 and HER2 in Gastric Carcinoma. Oncology 93, 387–394 (2017).

5. M. Song et al., PTEN loss increases PD-L1 protein expression and affects the correlation between PD-L1 expression and clinical parameters in colorectal cancer. PLoS One 8, e65821 (2013).

6. L. Wang et al., Clinical significance of B7-H1 and B7-1 expressions in pancreatic carcinoma. World J Surg 34, 1059–1065 (2010).

7. Y. Ye et al., Interaction of B7-H1 on intrahepatic cholangiocarcinoma cells with PD-1 on tumor-infiltrating T-cells as a mechanism of immune evasion. J Surg Oncol 100, 500–504 (2009).

8. M. Y. Zhang, Y. Y. Yang, X. H. Wang, X. F. Li, [Expression of Bcl-2, PD-L1 and its clinical significance in colorectal cancer]. Sichuan Da Xue Xue Bao Yi Xue Ban 43, 827–829, 859 (2012).

9. S. M. Ansell et al., PD-1 blockade with nivolumab in relapsed or refractory Hodgkin’s lymphoma. N Engl J Med 372, 311–319 (2015).

10. E. B. Garon et al., Pembrolizumab for the treatment of non-small-cell lung cancer. N Engl J Med 372, 2018–2028 (2015).

11. P. T. Nghiem et al., PD-1 Blockade with Pembrolizumab in Advanced Merkel-Cell Carcinoma. N Engl J Med 374, 2542–2552 (2016).

12. O. Hamid et al., Safety and tumor responses with lambrolizumab (anti-PD-1) in melanoma. N Engl J Med 369, 134–144 (2013).

13. S. L. Topalian et al., Safety, activity, and immune correlates of anti-PD-1 antibody in cancer. N Engl J Med 366, 2443–2454 (2012).

14. M. Ayers et al., IFN-gamma-related mRNA profile predicts clinical response to PD-1 blockade. J Clin Invest 127, 2930–2940 (2017).

15. T. Powles et al., MPDL3280A (anti-PD-L1) treatment leads to clinical activity in metastatic bladder cancer. Nature 515, 558–562 (2014).

16. N. J. Llosa et al., The vigorous immune microenvironment of microsatellite instable colon cancer is balanced by multiple counter-inhibitory checkpoints. Cancer Discov 5, 43–51 (2015).

17. D. T. Le et al., Mismatch repair deficiency predicts response of solid tumors to PD-1 blockade. Science 357, 409–413 (2017).

18. J. Grosso et al., Association of tumor PD-L1 expression and immune biomarkers with clinical activity in patients (pts) with advanced solid tumors treated with nivolumab (anti-PD-1; BMS-936558; ONO-4538). JTO 31, 3016–3016 (2013).

19. D. P. Carbone et al., First-Line Nivolumab in Stage IV or Recurrent Non-Small-Cell Lung Cancer. N Engl J Med 376, 2415–2426 (2017).

20. N. A. Rizvi et al., Cancer immunology. Mutational landscape determines sensitivity to PD-1 blockade in non-small cell lung cancer. Science 348, 124–128 (2015).

21. R. M. Samstein et al., Tumor mutational load predicts survival after immunotherapy across multiple cancer types. Nat Genet 51, 202–206 (2019).

22. A. C. Huang et al., T-cell invigoration to tumour burden ratio associated with anti-PD-1 response. Nature 545, 60–65 (2017).

23. C. S. Garris et al., Successful Anti-PD-1 Cancer Immunotherapy Requires T-cell-Dendritic Cell Crosstalk Involving the Cytokines IFN-gamma and IL-12. Immunity 49, 1148–1161 e1147 (2018).

24. T. R. Cottrell, J. M. Taube, PD-L1 and Emerging Biomarkers in Immune Checkpoint Blockade Therapy. Cancer journal (Sudbury, Mass.) 24, 41–46 (2018).

25. S. B. Coffelt et al., IL-17-producing gammadelta T-cells and neutrophils conspire to promote breast cancer metastasis. Nature 522, 345–348 (2015).

26. A. R. Abbas, K. Wolslegel, D. Seshasayee, Z. Modrusan, H. F. Clark, Deconvolution of blood microarray data identifies cellular activation patterns in systemic lupus erythematosus. PLoS One 4, e6098 (2009).

27. J. K. Neuenburg et al., T-cell activation and memory phenotypes in cerebrospinal fluid during HIV infection. J Acquir Immune Defic Syndr 39, 16–22 (2005).

28. S. Chen et al., Increased abundance of myeloid-derived suppressor cells and Th17 cells in peripheral blood of newly-diagnosed Parkinson’s disease patients. Neurosci Lett 648, 21–25 (2017).

29. A. R. Whitney et al., Individuality and variation in gene expression patterns in human blood. Proc Natl Acad Sci U S A 100, 1896–1901 (2003).

30. C. Krieg et al., High-dimensional single-cell analysis predicts response to anti-PD-1 immunotherapy. Nat Med 24, 144–153 (2018).

31. E. Azizi et al., Single-Cell Map of Diverse Immune Phenotypes in the Breast Tumor Microenvironment. Cell 174, 1293–1308 e1236 (2018).

32. M. Sade-Feldman et al., Defining T-cell States Associated with Response to Checkpoint Immunotherapy in Melanoma. Cell 175, 998–1013 e1020 (2018).

33. S. Hanzelmann, R. Castelo, J. Guinney, GSVA: gene set variation analysis for microarray and RNA-seq data. BMC Bioinformatics 14, 7 (2013).

34. A. Liberzon et al., The Molecular Signatures Database (MSigDB) hallmark gene set collection. Cell Syst 1, 417–425 (2015).

35. A. Liberzon et al., Molecular signatures database (MSigDB) 3.0. Bioinformatics 27, 1739–1740 (2011).

36. K. Schroder, P. J. Hertzog, T. Ravasi, D. A. Hume, Interferon-gamma: an overview of signals, mechanisms and functions. J Leukoc Biol 75, 163–189 (2004).

37. C. Shi, E. G. Pamer, Monocyte recruitment during infection and inflammation. Nat Rev Immunol 11, 762–774 (2011).

38. G. E. Mahlbacher, K. C. Reihmer, H. B. Frieboes, Mathematical modeling of tumor-immune cell interactions. J Theor Biol 469, 47–60 (2019).

39. J. Ozik et al., High-throughput cancer hypothesis testing with an integrated PhysiCell-EMEWS workflow. BMC Bioinformatics 19, 483 (2018).

40. M. Robertson-Tessi, A. El-Kareh, A. Goriely, A mathematical model of tumor-immune interactions. J Theor Biol 294, 56–73 (2012).

41. K. J. Mahasa, R. Ouifki, A. Eladdadi, L. Pillis, Mathematical model of tumor-immune surveillance. J Theor Biol 404, 312–330 (2016).

42. A. Carbo et al., Systems modeling of molecular mechanisms controlling cytokine-driven CD4+ T-cell differentiation and phenotype plasticity. PLoS Comput Biol 9, e1003027 (2013).

43. L. G. de Pillis, A. E. Radunskaya, C. L. Wiseman, A validated mathematical model of cell-mediated immune response to tumor growth. Cancer Res 65, 7950–7958 (2005).

44. H. C. Wei, J. L. Yu, C. Y. Hsu, Periodically Pulsed Immunotherapy in a Mathematical Model of Tumor, CD4(+) T-cells, and Antitumor Cytokine Interactions. Comput Math Methods Med 2017, 2906282 (2017).

45. R. Eftimie, J. L. Bramson, D. J. Earn, Interactions between the immune system and cancer: a brief review of non-spatial mathematical models. Bull Math Biol 73, 2–32 (2011).

46. J. L. Reading, S. A. Quezada, Too Much of a Good Thing? Chronic IFN Fuels Resistance to Cancer Immunotherapy. Immunity 45, 1181–1183 (2016).

47. M. Dosset et al., PD-1/PD-L1 pathway: an adaptive immune resistance mechanism to immunogenic chemotherapy in colorectal cancer. Oncoimmunology 7, e1433981 (2018).

48. I. Peguillet et al., High numbers of differentiated effector CD4 T-cells are found in patients with cancer and correlate with clinical response after neoadjuvant therapy of breast cancer. Cancer Res 74, 2204–2216 (2014).

49. R. Verma et al., Lymphocyte depletion and repopulation after chemotherapy for primary breast cancer. Breast Cancer Res 18, 10 (2016).

50. M. Palma et al., T-cells in chronic lymphocytic leukemia display dysregulated expression of immune checkpoints and activation markers. Haematologica 102, 562–572 (2017).

51. P. Yang, J. Ma, X. Yang, W. Li, Peripheral CD4+ naive/memory ratio is an independent predictor of survival in non-small cell lung cancer. Oncotarget 8, 83650–83659 (2017).

52. A. Dobin et al., STAR: ultrafast universal RNA-seq aligner. Bioinformatics 29, 15–21 (2013).

53. A. Butler, P. Hoffman, P. Smibert, E. Papalexi, R. Satija, Integrating single-cell transcriptomic data across different conditions, technologies, and species. Nat Biotechnol 36, 411–420 (2018).

54. I. Tirosh et al., Dissecting the multicellular ecosystem of metastatic melanoma by single-cell RNA-seq. Science 352, 189–196 (2016).

55. A. M. Newman et al., Robust enumeration of cell subsets from tissue expression profiles. Nat Methods 12, 453–457 (2015).

56. M. De Simone et al., Transcriptional Landscape of Human Tissue Lymphocytes Unveils Uniqueness of Tumor-Infiltrating T Regulatory Cells. Immunity 45, 1135–1147 (2016).

57. J. H. L. McInnes, J. Melville, UMAP: Uniform Manifold Approximation and Projection for Dimension Reduction. arXiv e-prints, (2018).

58. G. Finak et al., MAST: a flexible statistical framework for assessing transcriptional changes and characterizing heterogeneity in single-cell RNA sequencing data. Genome Biol 16, 278 (2015).

59. B. Bischl et al., mlr: Machine Learning in R. JMLR 17, 1–5 (2016).

60. K. Moon et al., Manifold Learning-based Methods for Analyzing Single-Cell RNA-Sequencing Data. Current Opinion in Systems Biology 7, (2017).

61. D. van Dijk et al., Recovering Gene Interactions from Single-Cell Data Using Data Diffusion. Cell 174, 716–729 e727 (2018).

62. K. Moon et al., Visualizing Structure and Transitions for Biological Data Exploration. SSRN Electronic Journal, (2018).

63. D. Risso, F. Perraudeau, S. Gribkova, S. Dudoit, J. P. Vert, A general and flexible method for signal extraction from single-cell RNA-seq data. Nat Commun 9, 284 (2018).

64. N. Lawrence, Probabilistic non-liner component analysis with Gaussian process latent variable models. JMLR 6, 1783–1816 (2005).

65. B. Carpenter et al., Stan: A Probabilistic Programming Language. 2017 76, 32 (2017).

