## Supporting information for "Circulating immune cell phenotype dynamics reflect the strength of tumor-immune cell interactions in patients during immunotherapy"

**Classification**

BIOLOGICAL SCIENCES: Systems Biology

**Affiliations**

**^1^** Department of Medical Oncology & Therapeutics Research, City of Hope National Medical Center, 1500 East Duarte Road, Duarte, CA.

**^2^** Department of Mathematics, University of Utah 155 S 1400 E RM 233 Salt Lake City, UT.

**^3^** College of Pharmacy, University of Minnesota, Duluth, MN, USA.

**^4^** Department of Integrative Biology and Pharmacology, School of Medicine, School of Biomedical Informatics, UT Health Sciences Center at Houston, Houston, TX, USA.

**^5^** Department of Oncological Sciences, School of Medicine, University of Utah, 2000 Circle of Hope Drive, Salt Lake City, UT 84112, USA.

**^6^** Department of Pharmacology and Toxicology, College of Pharmacy, University of Utah, 30 South 2000 East, Salt Lake City, UT 84112, USA.

**^7^** Department of Pathology and Laboratory Medicine, David Geffen School of Medicine, University of California Los Angeles, CA 90095, USA.

**^8^** Translational Oncology Research & Drug Discovery, Translational Genomics Research Institute (TGen), 445 N. Fifth Street, Phoenix, AZ 85004, USA.

† These authors contributed equally to this work

**Corresponding Authors**:

Andrea Bild, Ph.D. Professor,

Department of Medical Oncology and Therapeutics

City of Hope Comprehensive Cancer Institute

1218 S Fifth Ave, Monrovia, CA 91016

Sunil Sharma, MD, FACP, MBA

Deputy Director, Clinical Sciences Professor and Division Director

Translational Genomics Research Institute (TGen)

Chief, Translational Oncology Research & Drug Discovery

Honor Health Research Institute

Professor of Medicine, City of Hope

**This PDF file includes:**

Supplementary text

Figures S1 to S16

Tables S1 to S3

**Other supplementary materials for this manuscript include the following:**

Dataset S1 to S4

**Supplementary text**

**Supplementary methods**

**Study overview**

The objective of this study was to analyze gene expression profiles and number of circulating immune cells of patients treated with a sequenced combination of chemotherapy (mFOLFOX6) and immunotherapy (anti-PD-1) at different time points (without treatment, 1 month after chemotherapy and 1 month after chemo and immunotherapy). Data collection began on 1/28/2015 and was completed on 5/22/2018.

**Immune cell annotation verification**

*Machine learning method*

PD-1 therapy related immune cell scRNAseq samples (N = 69745) were used to train and evaluate the classifiers. scRNAseq raw counts for PBMCs were downloaded from GEO accession GSE114727 (*1*) as a test data set. TPM values for tumor-infiltrating immune cells downloaded from GSE120575 (*2*) were used as a second test data set. In each PD-1 dataset, 500 samples were randomly selected from each cluster, except for T6. For clusters with less than 1000 cells, oversampling was performed for 50% of the cells in the cluster.  In total, 17,000 cells were used as a training set. The rest of the samples were held out for validation. We repeated this procedure and generated 10 different training and 10 corresponding validation sets.

We ran Seurat on the training dataset to obtain the cluster-associated markers (*3*). Only positive markers were selected. For efficiency, only genes that were detected in a minimum fraction of 0.25 in either of the two populations (min.pct=0.25) and whose average expression was larger than 0.25 between clusters were selected (thresh.use=0.25). We ranked the candidate markers using adjusted p-values. We selected the top 20 markers per cluster (410 unique genes) as features in the machine learning process. We performed a multi-class classification using MLR with the RandomForest classifier (*4*).

*Shared markers method*

We downloaded the lists of markers from Azizi et al. and Sade-Feldman et al. (*1, 2*). The markers in Azizi et al. were ranked by z-score and only those with absolute z-score greater than 1.95 were kept. Then we calculated the number of shared marker genes between public data sets and top 20 markers for each cluster in our study. We used a network structure to visualize the similarity and hierarchy of our clusters and the ones in a breast cancer study. The rationale of inferring that cluster B belongs to A is that: (i) A and B share a high number of markers, and (ii) A can pair more clusters than B. The network visualization workflow is shown in Fig S4.

**Quantifying immune cell phenotypes**

We measured the phenotypic variation of cell states within cell types, using affinity-based pseudotime reconstruction (*5, 6*). It also allowed complex nonlinear gene expression changes to be captured, despite the sparsity of transcripts, the presence of amplification noise and normalization induced artifacts that is present in all single cell datasets (*7*). We filtered out genes that had zero expression in more than 99% of cells. The first 150 principle components of this gene expression profile were then calculated and used to create a dissimilarity matrix (Euclidean distance) of gene expression between cells. Nearest neighbor distances were calculated for the 100 most similar cells and an exponential kernel was used to produce the local affinity matrix, which was normalized to give the Markov transition matrix between cells. The kernel decay rate (α) parameter was set to 10, to capture the local and global relatedness between cells (Parameter sensitivity was low but below 5 the global structure was lost and above 30 local structure was lost). Log transition probabilities were calculated and a reduce dimension embedding was created to visualize the first 5 major axes of phenotypic variation.

**Estimating tumor burden time courses using Gaussian process models**

Gaussian processes latent variable models were fitted to each patients’ tumor time course data (Cancer antigen measurements and CT scan results). A Bayesian priors of nonlinear functions was constructed to describe a wide range of different kinds of fluctuation in tumor burden that could have occurred. Then, using the Markov chain Monte Carlo, we infer a posterior that represents our belief about the fluctuations that are likely to have occurred, conditioned on observed data. Specifically, a multidimensional Gaussian,$F=\left( f_{1} \ldots f_{x} \right)\sim N\left( \mu,\Sigma\right), \{indices i=1\ldots x\}$, was defined, with each of $n$ dimensions describing the tumor burden at one of the time points at which either RECIST CT or blood biomarker data was available. The nonlinear function describing the tumor burden over time ($f\left( t \right)$) was then described as a Gaussian process: $f\left( t \right)\sim GP\left( m\left( x \right), k\left( x,x^{'} \right) \right), \left\{ indices x \right\},$where$m\left( x \right)$is an $n$-vector and$k\left( x,x^{'} \right)$ is an $n$x$n$ covariance matrix. The covariance of tumor burden at different time points was calculated, using a squared exponential covariance function:

$k{(x)}_{i,j}= \eta^{2}\exp\left( -\rho^{2}\sum_{d=1}^{D} \left( x_{i,d} -x_{j,d} \right)^{2} \right)+ \delta_{i,j}\sigma^{2}$.

Covariance function hyper-parameters ($\eta,$ $\rho$) were estimated. To ensure a positive definite covariance matrix, $\sigma^{2}$ was added to the diagonal elements, using $\delta$ as a Kronecker delta function with value 1 if $i=j$, but 0 otherwise. The summation of squared Euclidean distances in time results in smooth changes in tumor burden over time.

RECIST1.1 assessments of the volume of tracked lesions within a given patient ($R_{P}$) was assumed to be a lognormally distributed measurement of a fixed fraction of a patient’s overall tumor burden: $R_{P}(t)\sim LogNormal(f_{P}\left( t \right),\sigma_{R_{P}} )$. The two blood biomarkers were assumed to be produced by cancer cells at a constant rate, $\alpha$, degrade at rate $\lambda$ and to have a baseline production level within the healthy body of rate $r$. The dynamics of blood antigens can then be described as: $\frac{dM_{x}}{dt}=r+\alpha f_{P}\left( t \right)- \lambda M_{x}$. Applying steady state assumptions, the blood biomarkers were also expected to be lognormally distributed measurements such that: ${M_{x}}_{P}\left( t \right)\sim LogNormal\left( {\beta_{x}}_{P} f_{P}\left( t \right)+ {c_{x}}_{P} , \sigma_{{M_{x}}_{P}} \right)$. Bayesian sampling of these probabilistic models yielded overall time course estimates of tumor burden during the trial. Concurrent measurements of immune cell abundance and therapeutic dosages (immunotherapy and chemotherapy) were also curated.

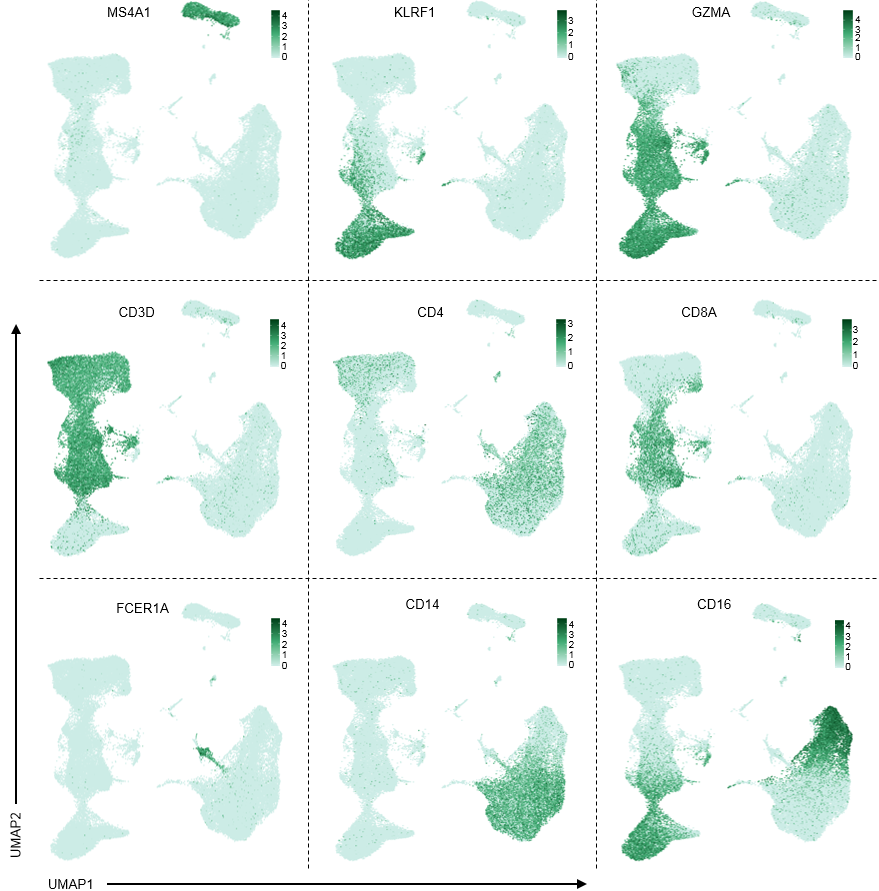

Fig. S1: UMAP key gene expression markers of major PBMC populations.

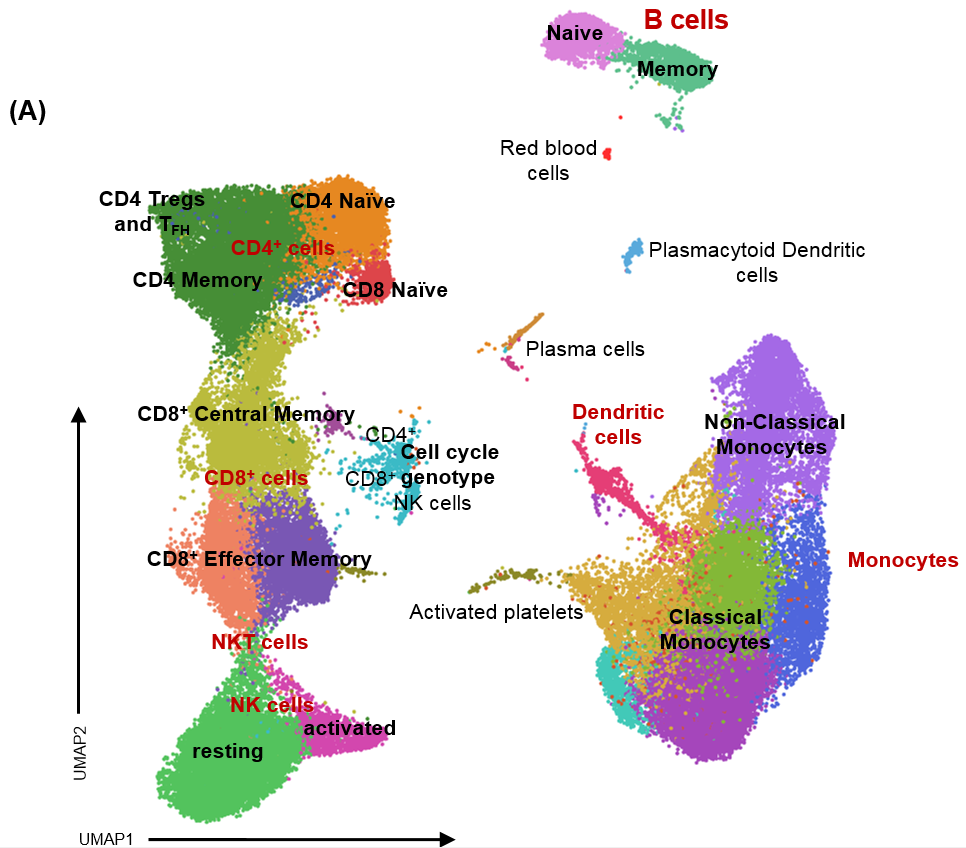

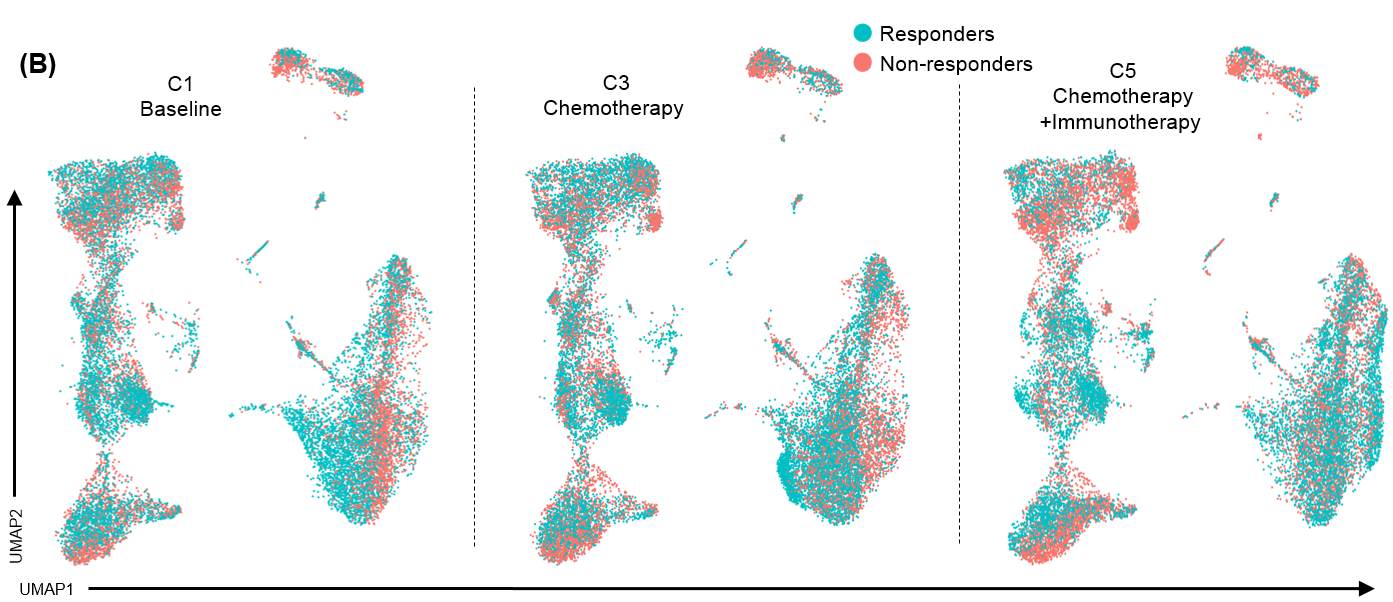

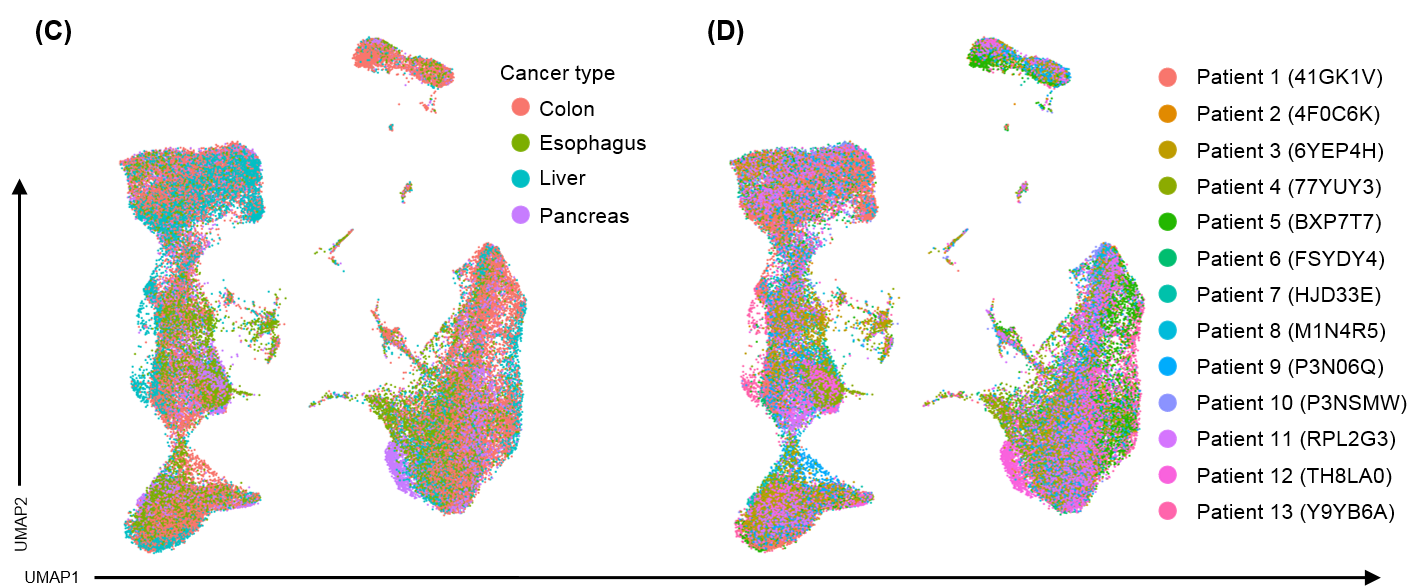

Fig. S2: UMAP gene expression shows no batch effect **(A)** Uniform Manifold Approximation and Projection (UMAP) of the scRNA seq data of all patients and all time point analyzed. Major PBMC clusters are labeled in red while subclusters or minor PBMC clusters are labeled in black. **(B)** UMAP gene expression distribution per time point of treatment. C stands for cycle, 1 cycle is 14 days, C1= baseline, C3= chemotherapy mFOLFOX6 regimen, C5 and 8= Chemotherapy + anti PD-1 immunotherapy. **(C)** UMAP gene expression distribution per patient analyzed. **(D)** UMAP gene expression distribution per cancer analyzed.

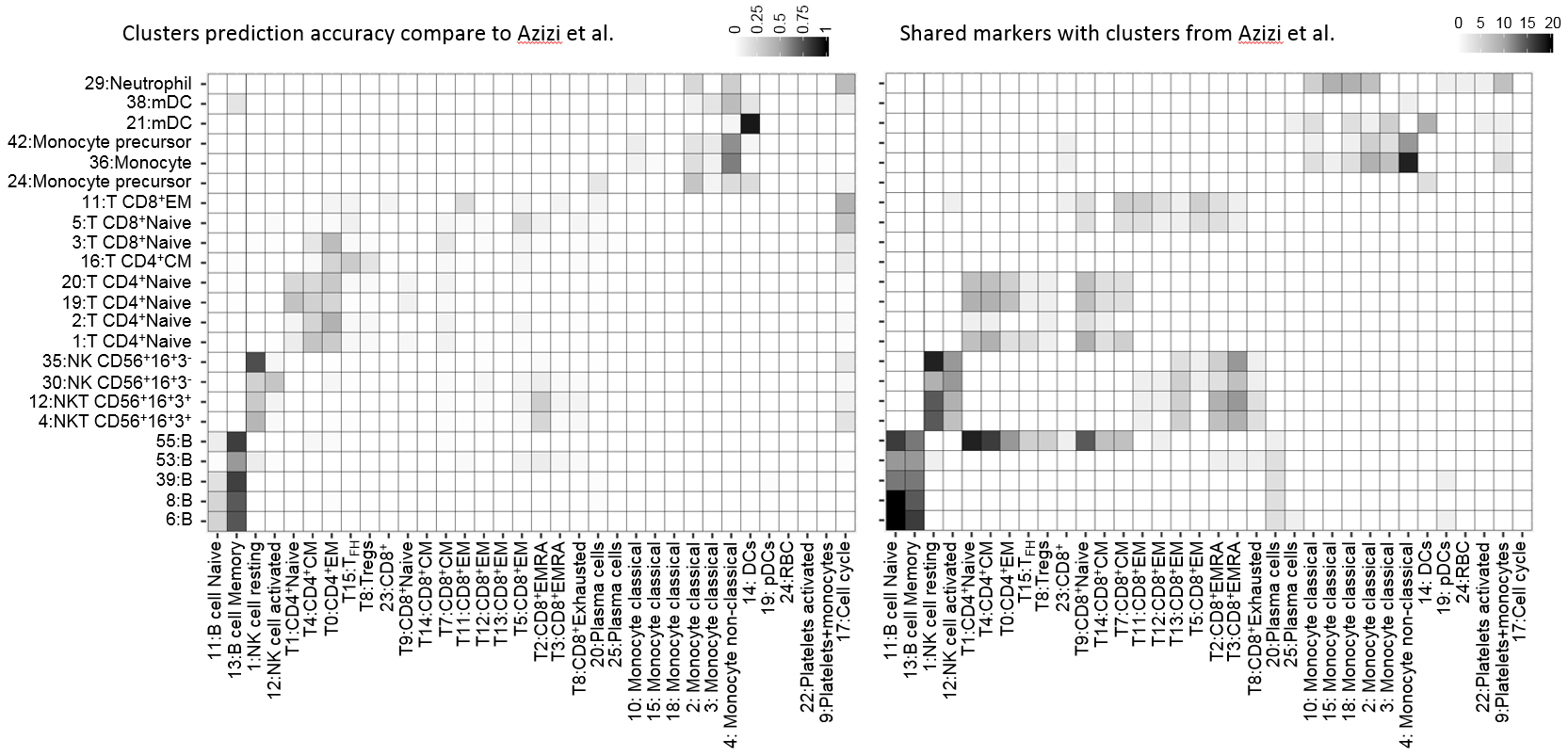

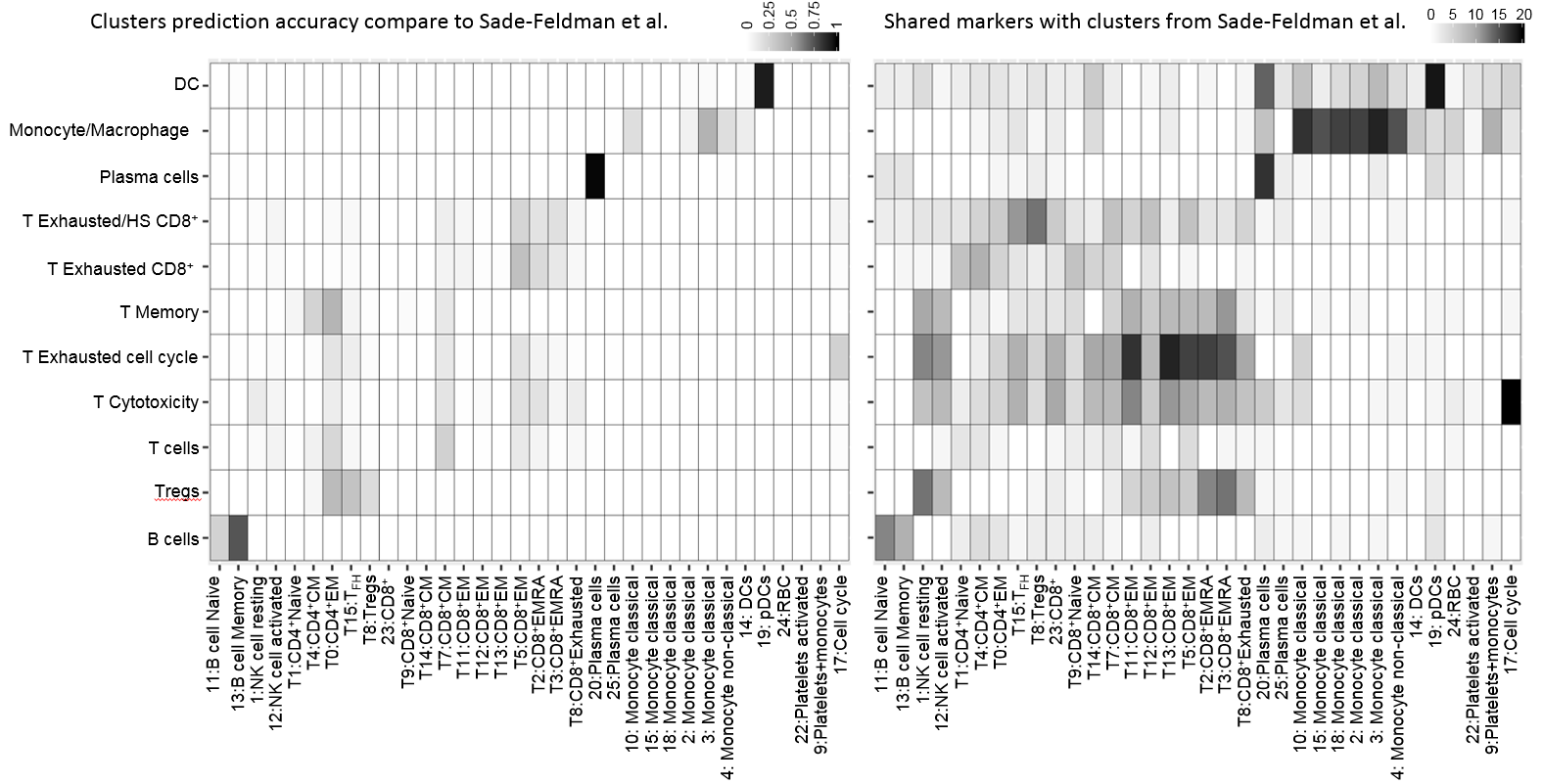

Fig. S3: UMAP labeled clusters are similar to public datasets. Left panels (machine learning prediction): the distribution of immune cells in public datasets predicted to PD-1 clusters by Random Forest learner using our cohort as a training set. Right panels (Shared marker genes): the number of shared genes between public datasets and our cohort clusters. Top panels corresponds to clusters published by Azizi et al.(*30*) while the bottom panels correspond to clusters published by Sade-Feldman et al.(*29*)

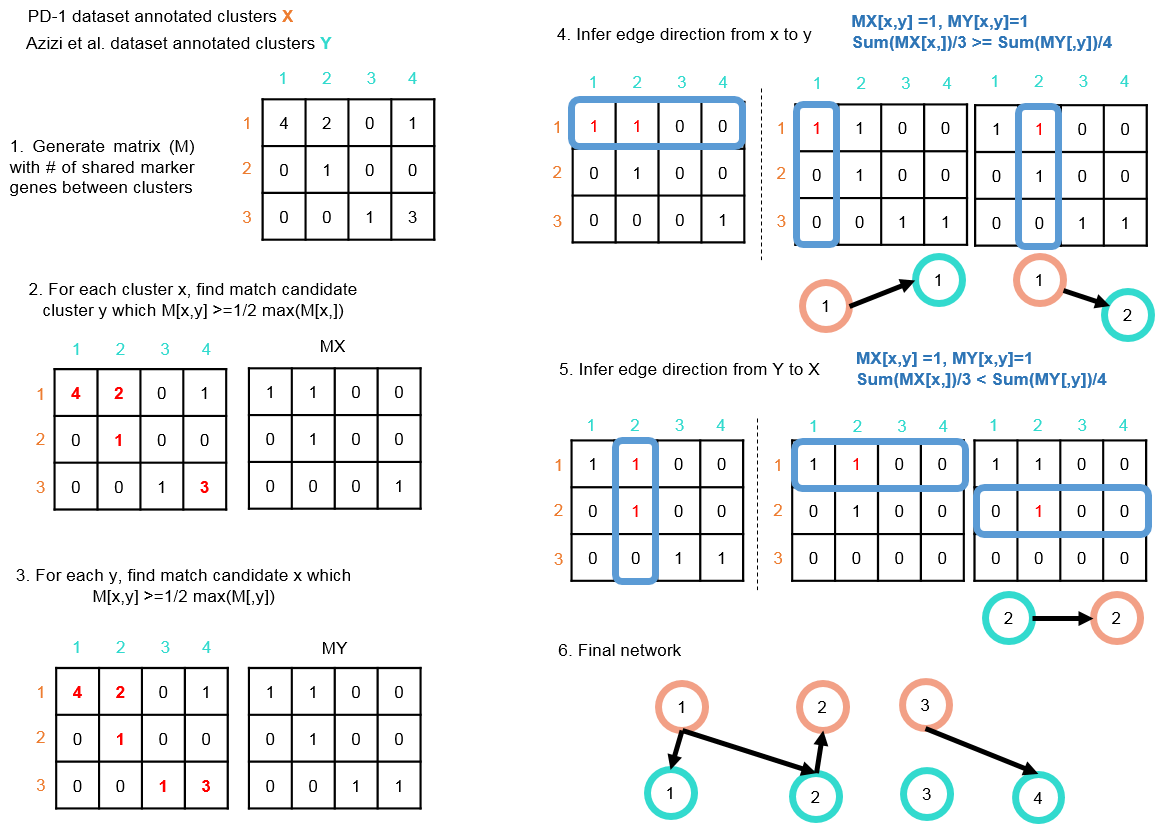

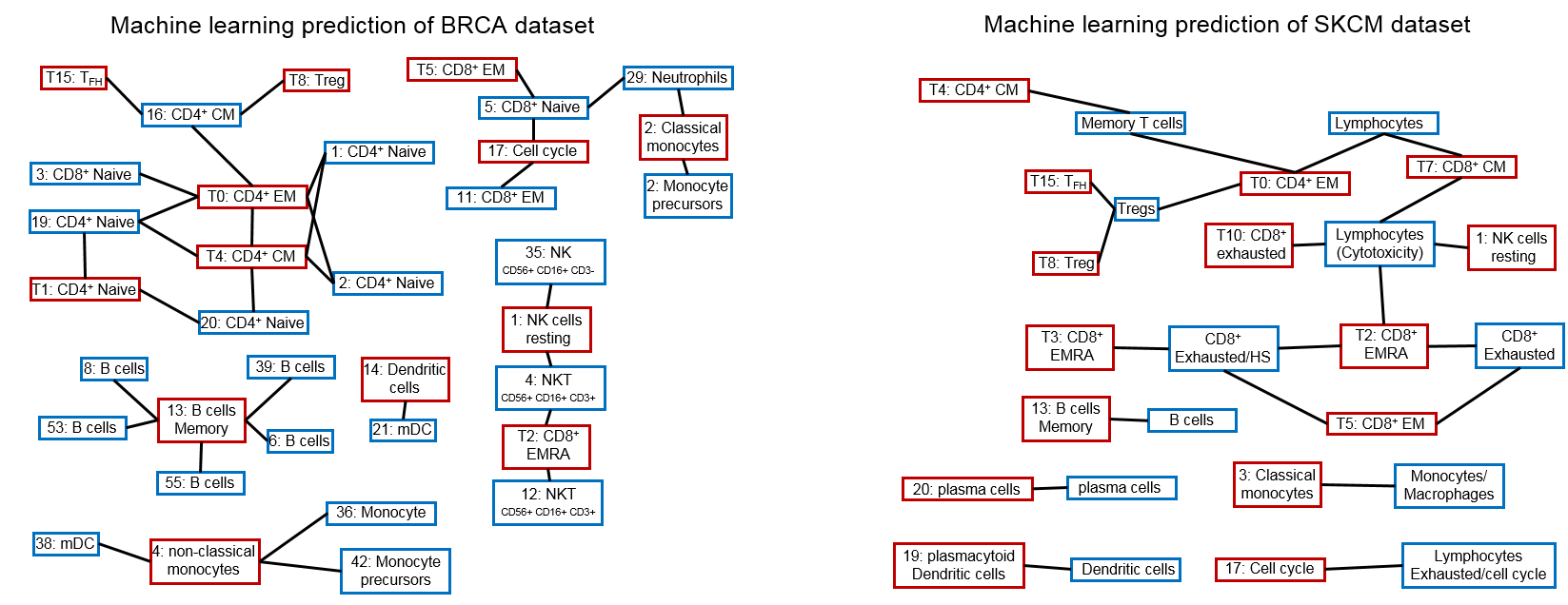

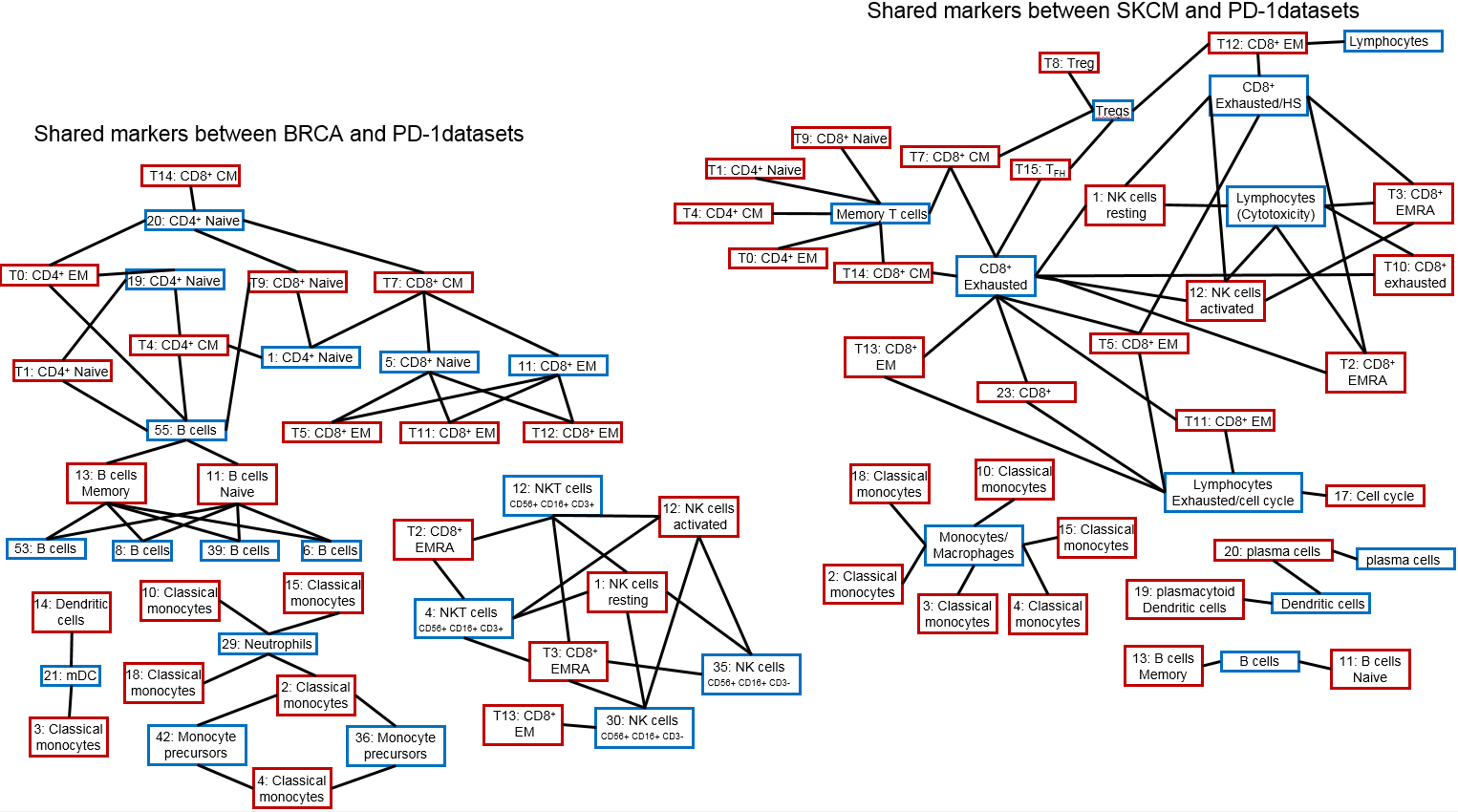

Fig. S4: Connections between our annotated clusters and public dataset clusters. Top panel (machine learning prediction): Connections of immune cells in public datasets predicted to PD-1 clusters by Random Forest learner using our cohort as a training set. Bottom panel (Shared marker genes): Connections of the number of shared genes between public datasets and our cohort clusters. Right panel corresponds to cluster published by Azizi et al.(30) while the right panel corresponds to cluster published by Sade-Feldman et al.(*29*).

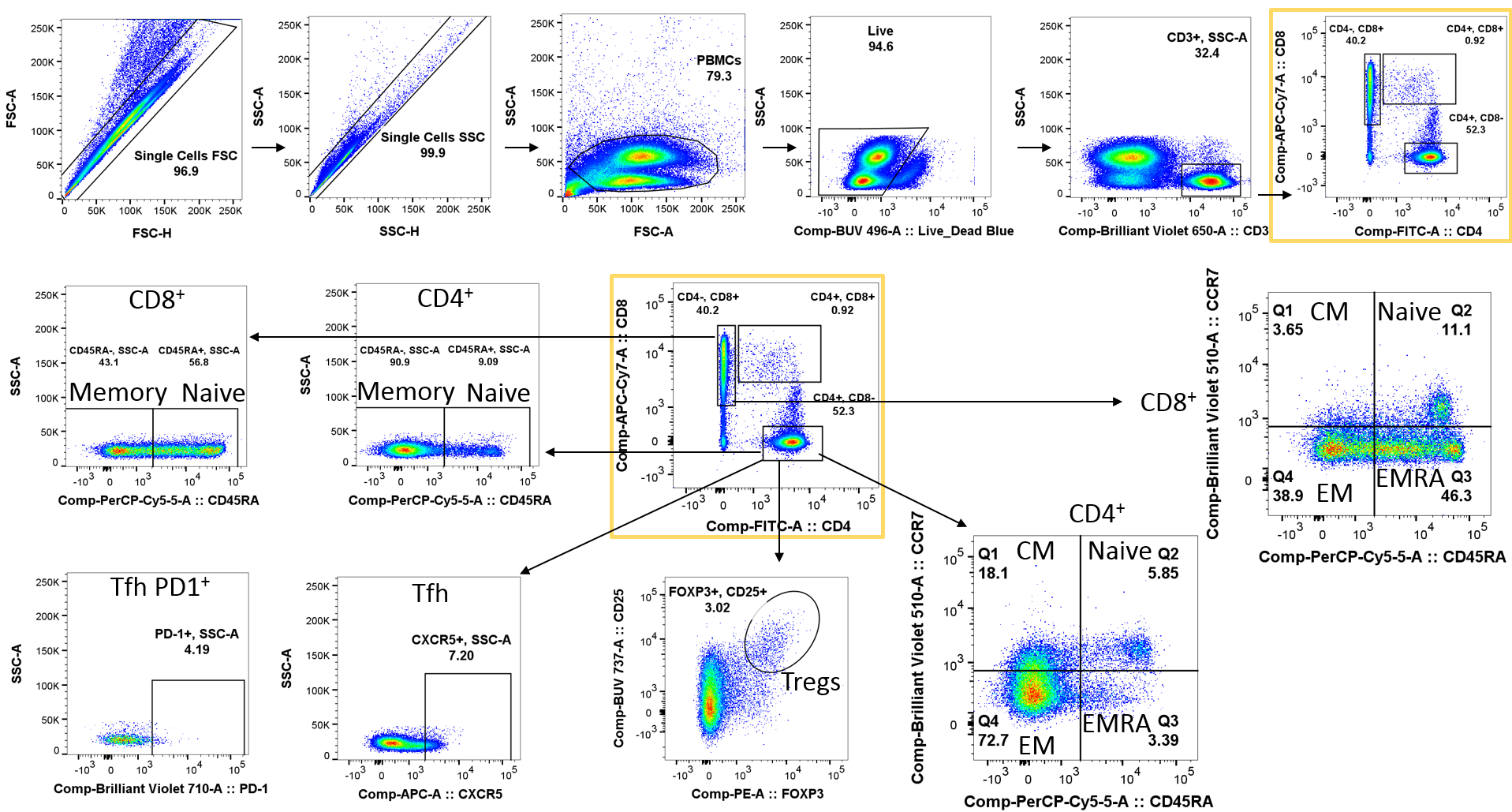

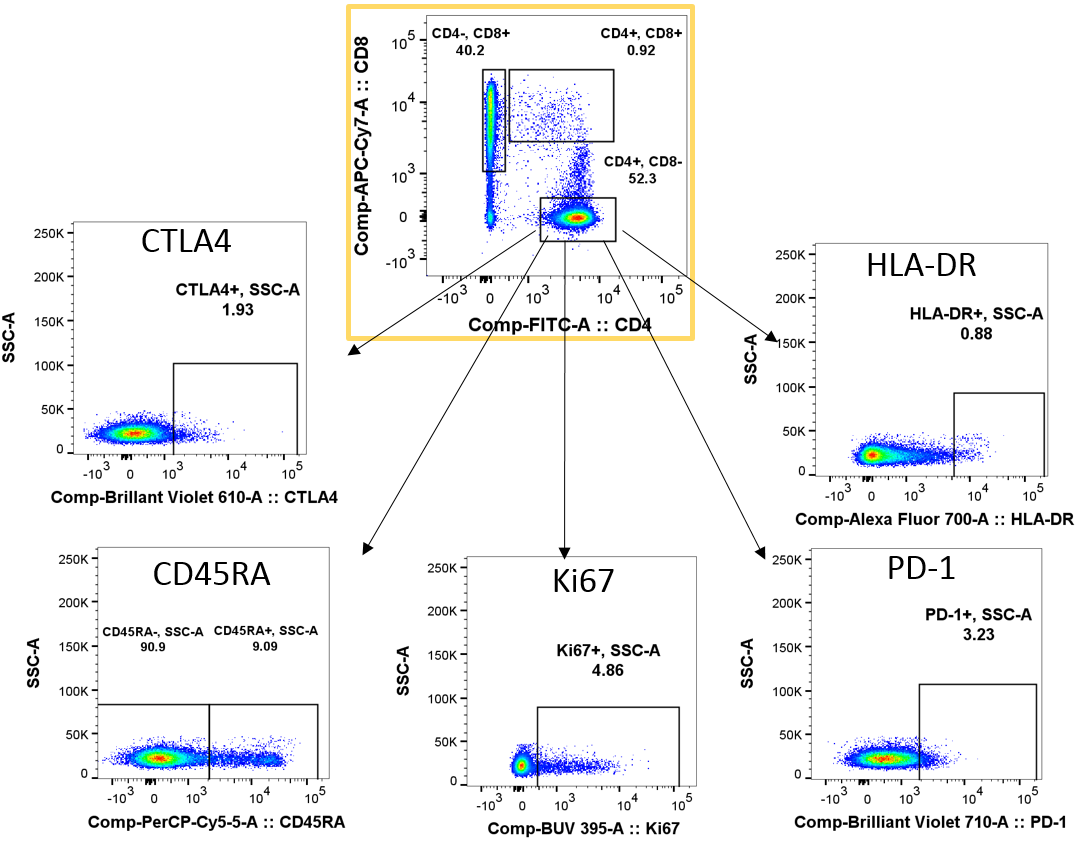

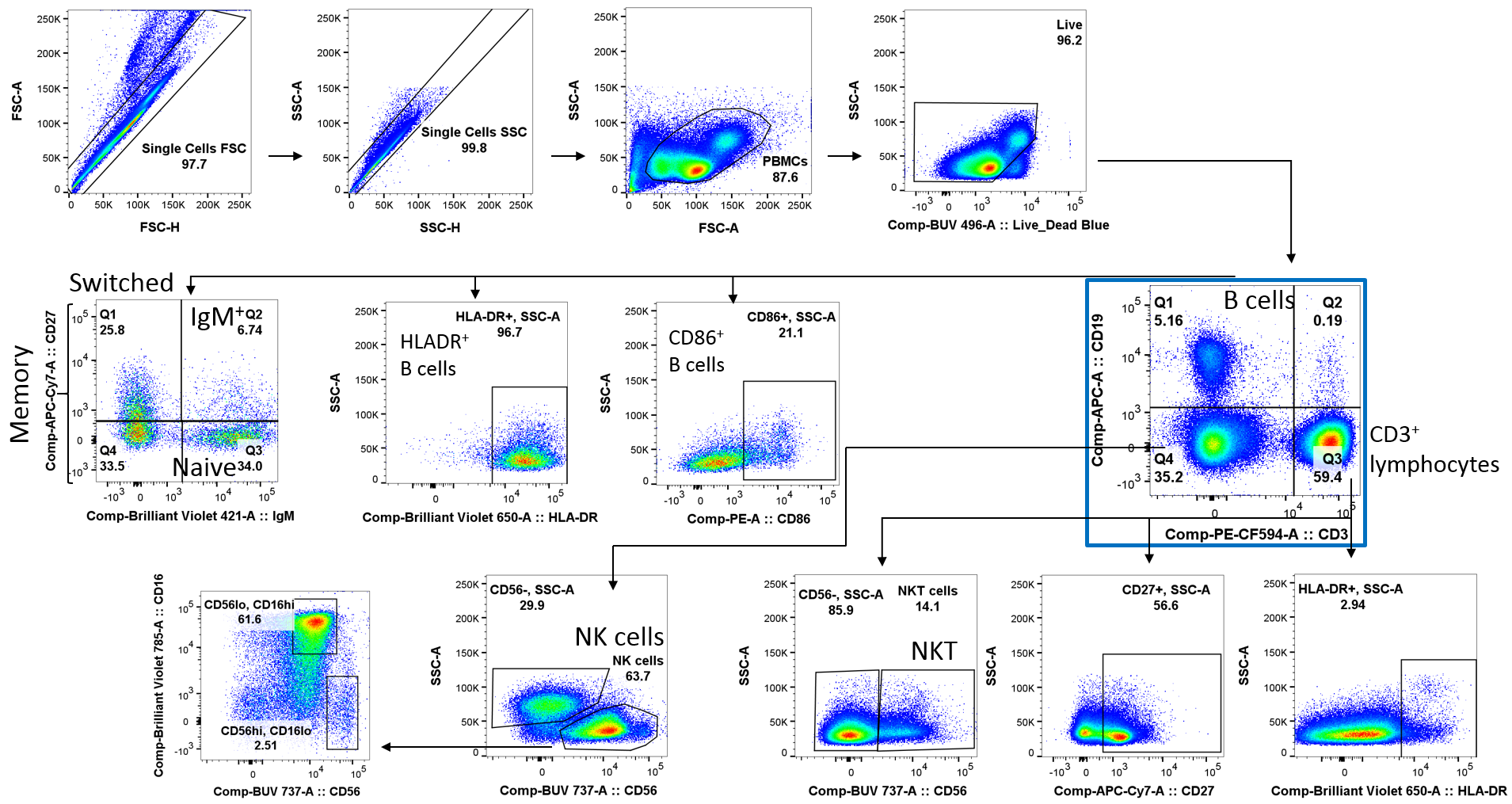

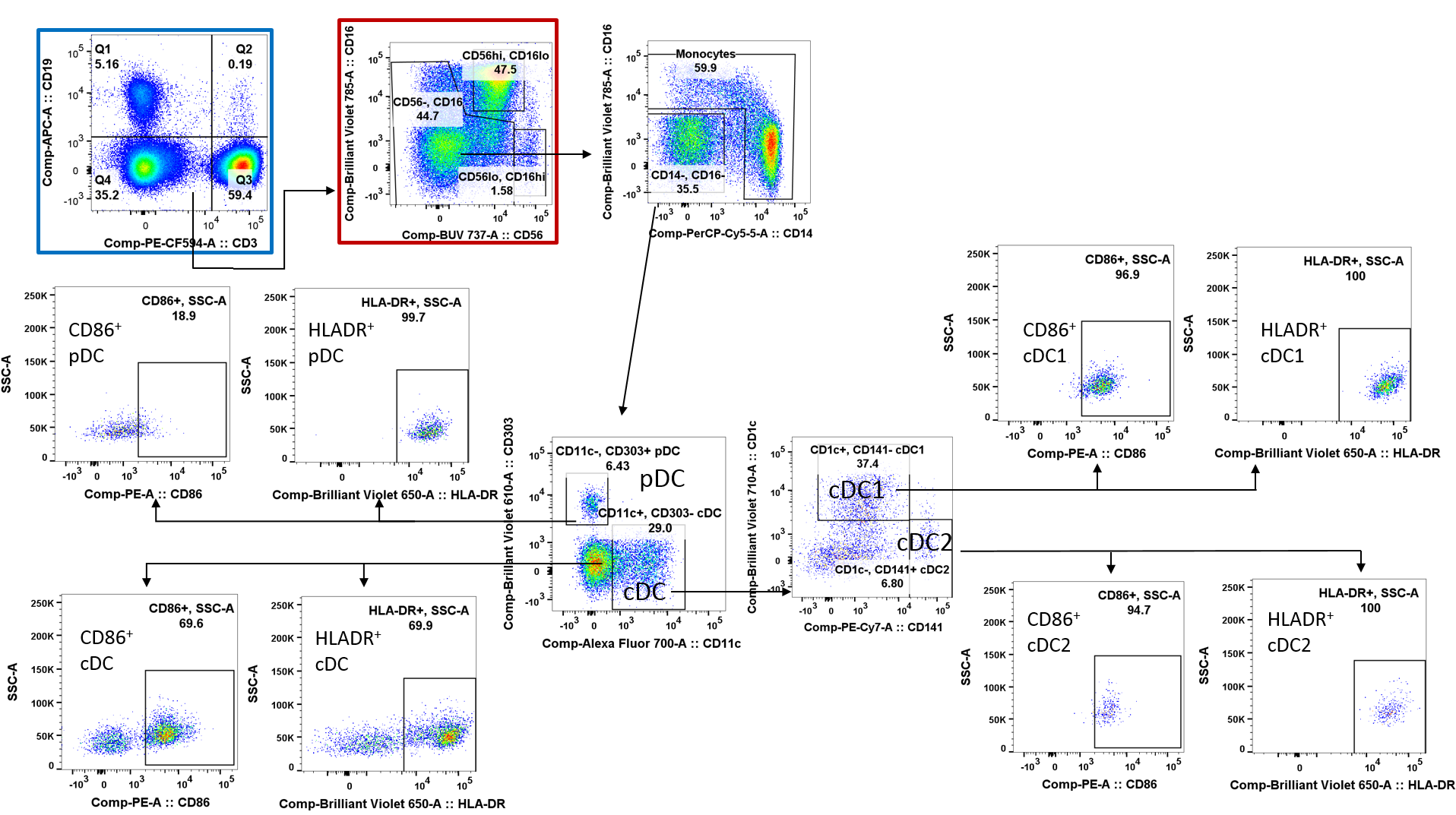

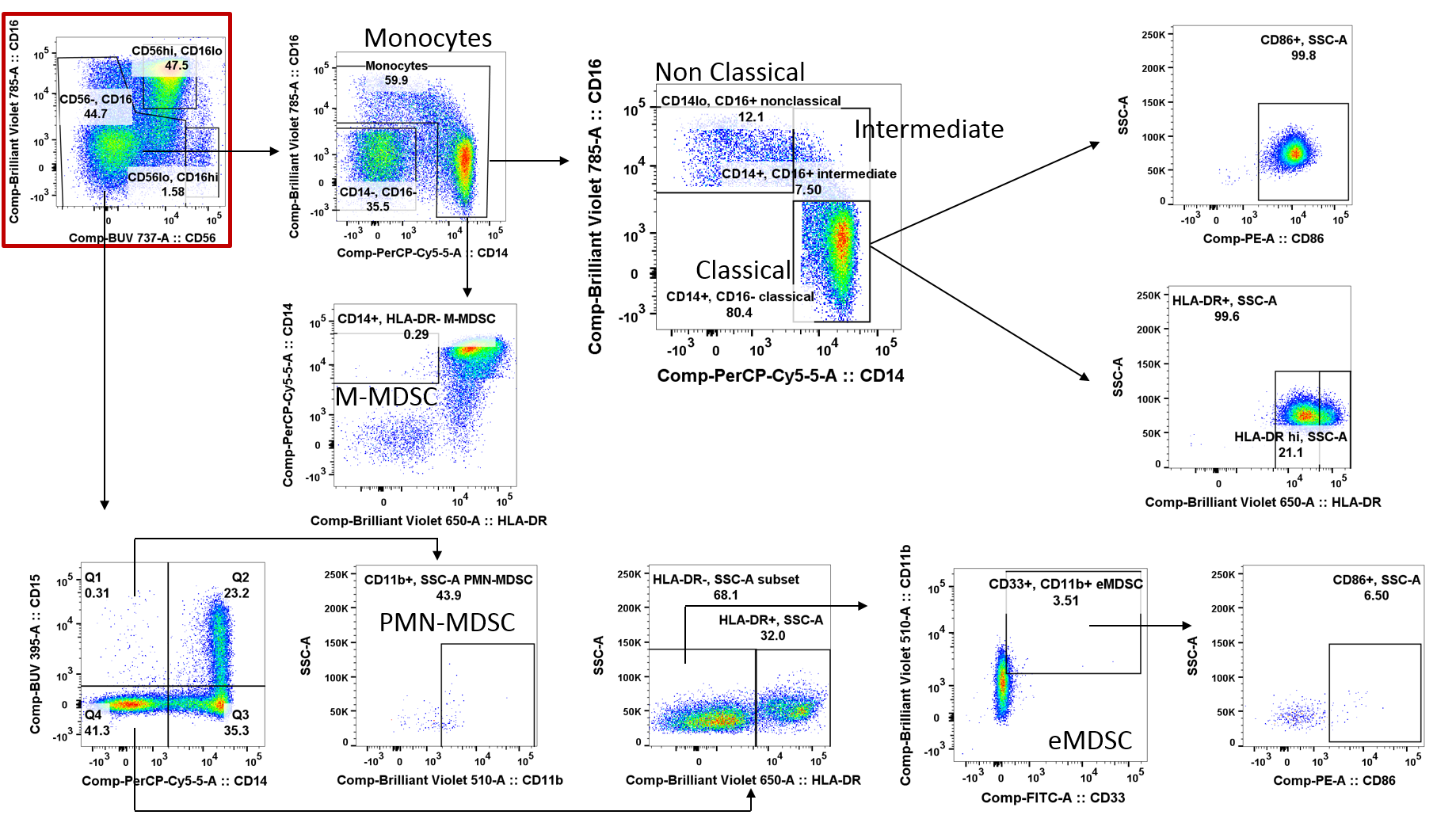

Fig. S5: Gating strategy used to analyze PBMCs.

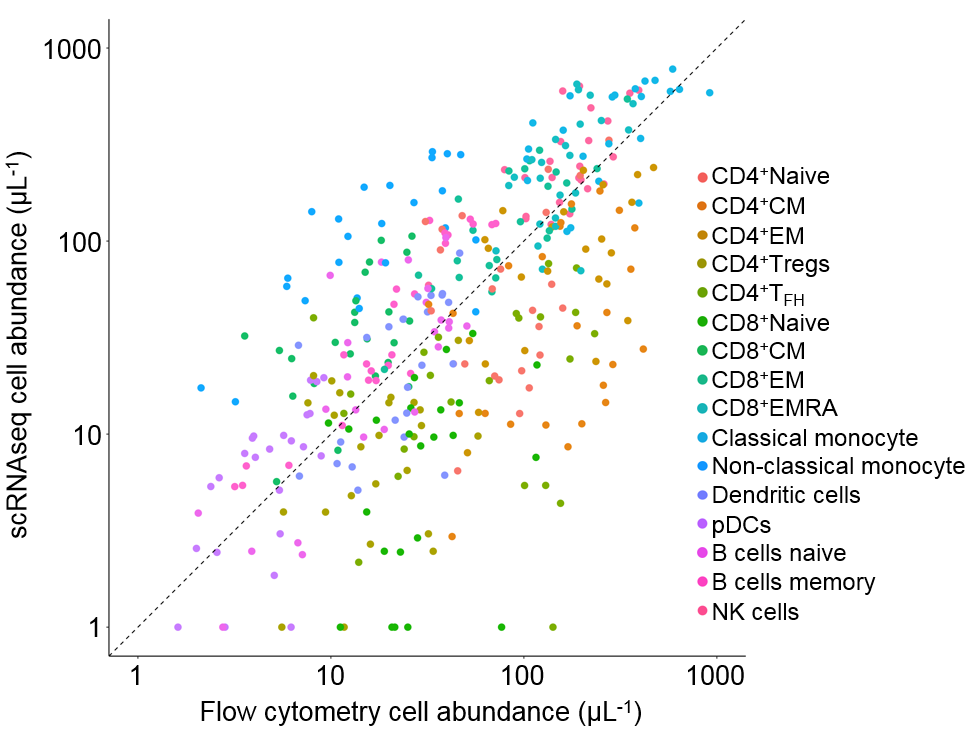

Fig. S6: Comparison of immune cell subtype abundances, as estimates using scRNA-seq or flow cytometry. Abundances of each cell subtype (colored points) were obtained scaling cell proportions measured by each method by the observed PBMC abundance in the blood sample. Cell subtype abundance was calculated for each patient, at each time point. The diagonal dashed line shows the 1:1 correspondence between the approaches.

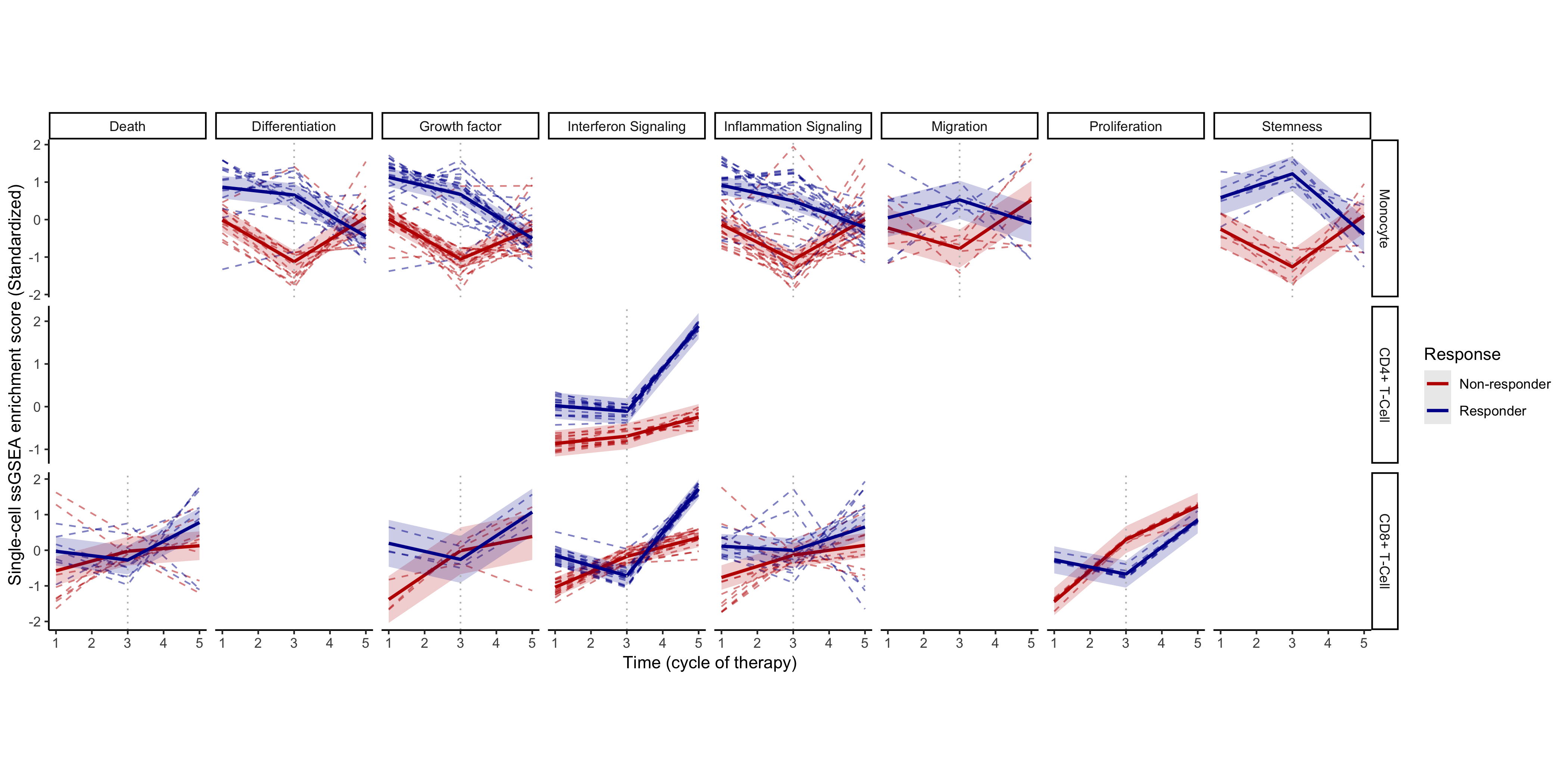
Fig. S7: Overview of differential temporal trends in monocyte and T cell pathway expression between responders and non-responders. Response (color) related differences in pathway activity was identified relating to: cell death, differentiation, growth factor production, interferon, and inflammatory signaling, migration and stemness. These pathway changes are visualized when present for monocytes, CD4^+^, and CD8^+^T cells. Trends of pathways exhibiting differential expression patterns in responders and non-responders are indicated by dashed lines. Overall trends of pathways within each cellular process (solid lines) and variation (shaded regions) are overlaid.

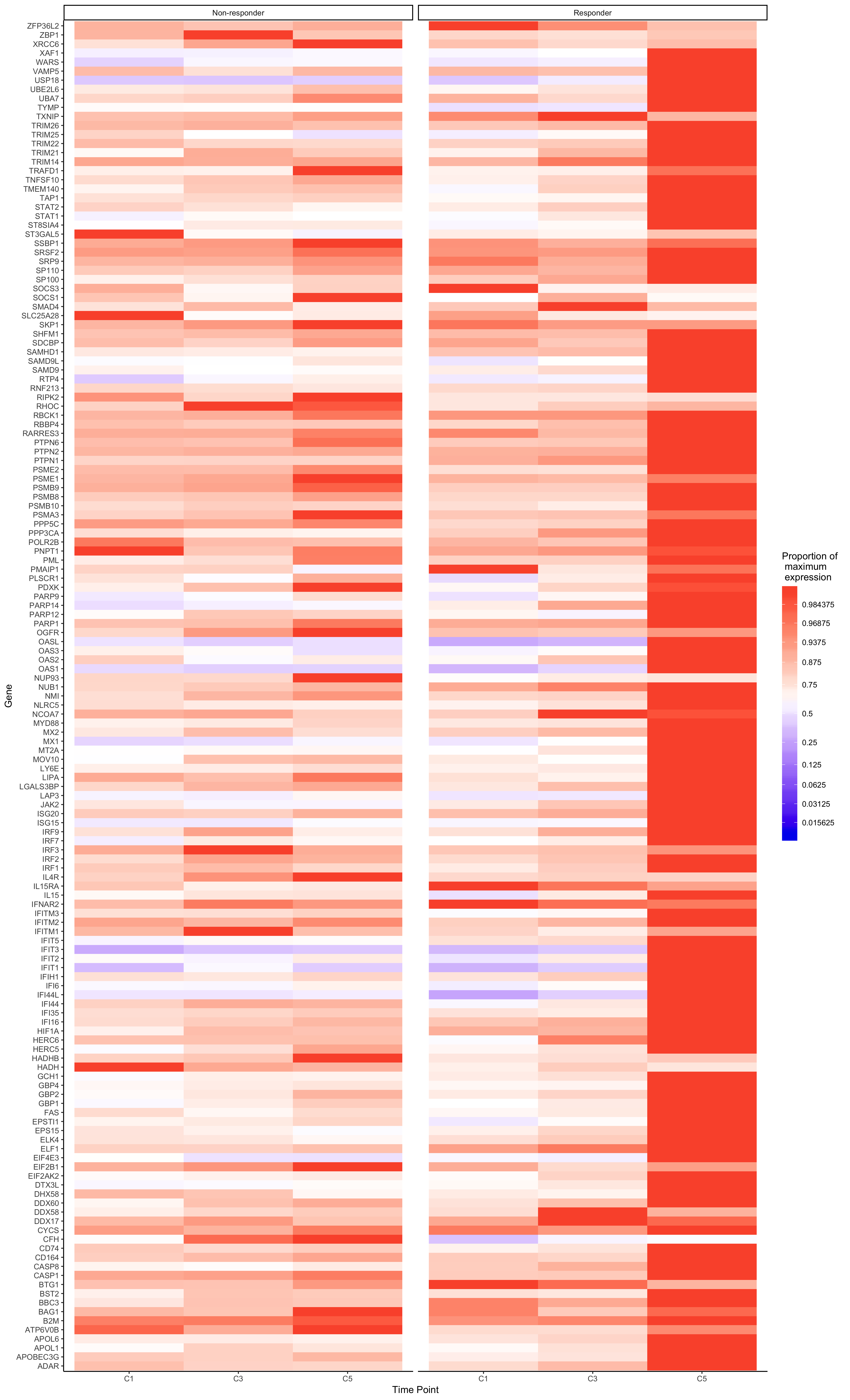

CD4 IFN signalling

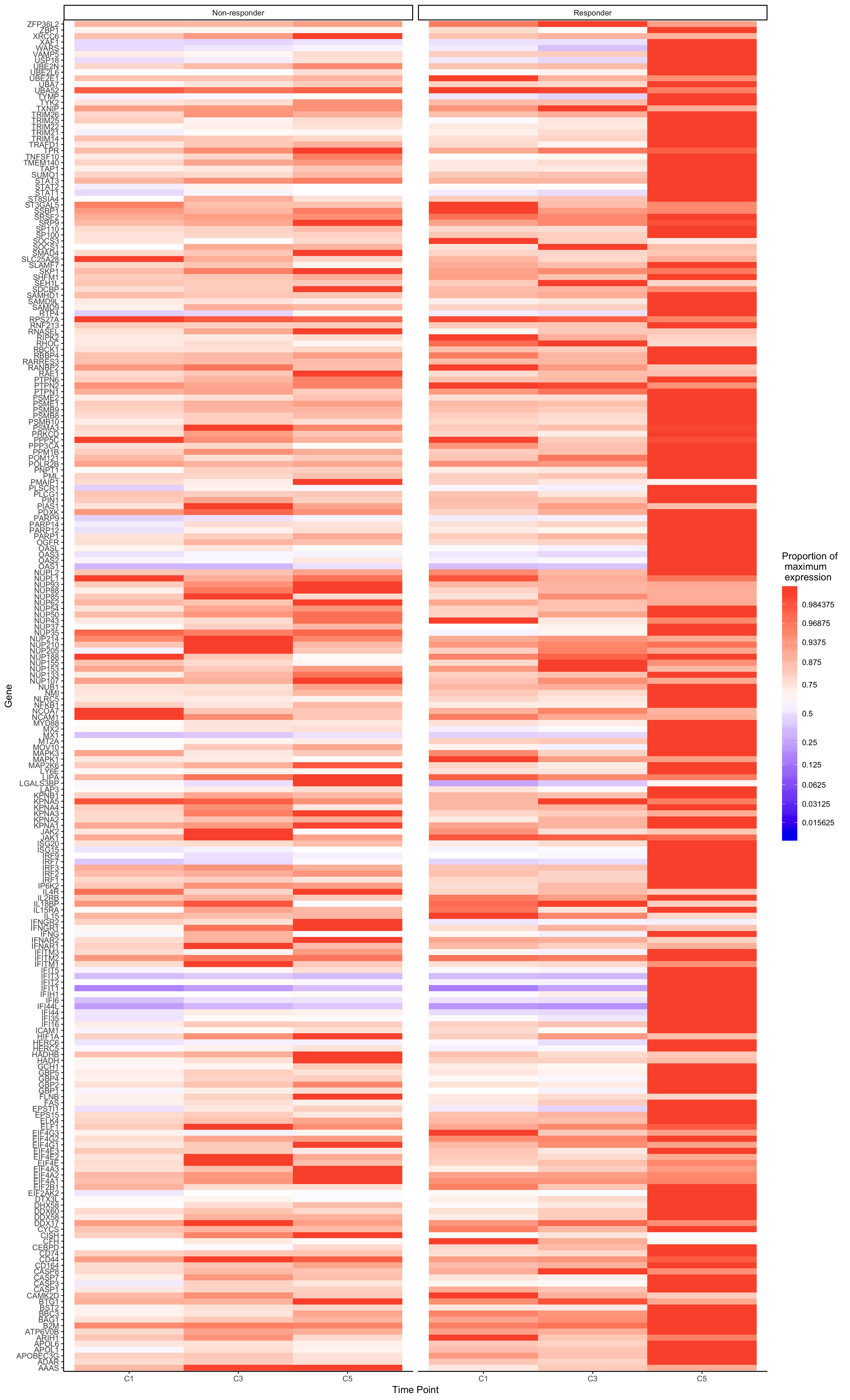

CD8 IFN signalling

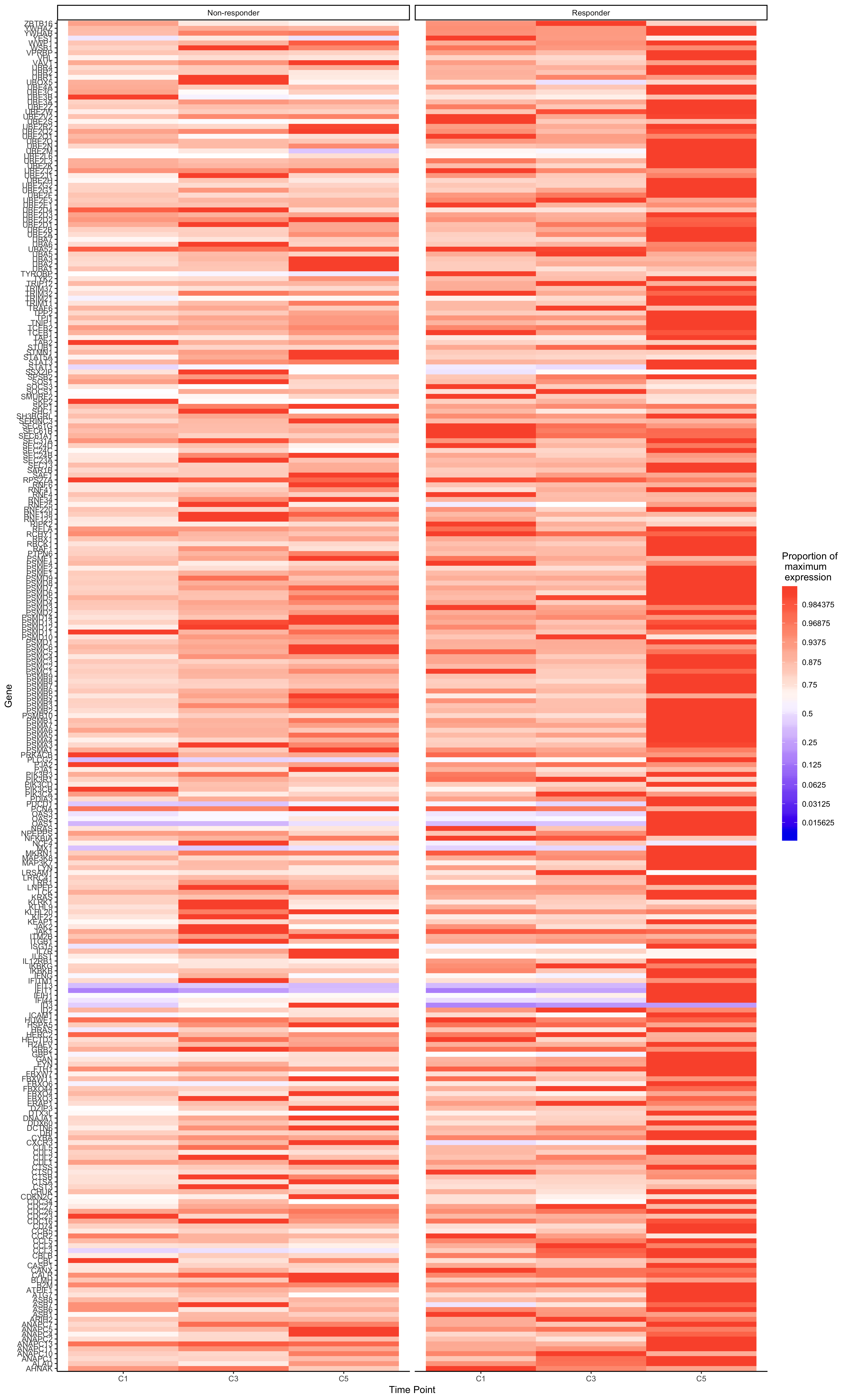

CD8 Inflammation

Fig. S8: Responders CD4^+^ and CD8^+^ T cells show upregulation of IFN gene signature pathways following the addition of anti-PD-1 immunotherapy. Heatmaps of changes in IFN (interferon) pathway-related gene expression of responders and non-responders CD4^+^ and CD8^+^ T cells over time. Gene expression is displayed as the proportion of the maximum level of each gene. Expression compared at time points C1 (baseline), C3 (start of chemotherapy mFOLFOX6 regimen) and C5 (Two cycles of anti-PD-1 immunotherapy in addition to chemotherapy).

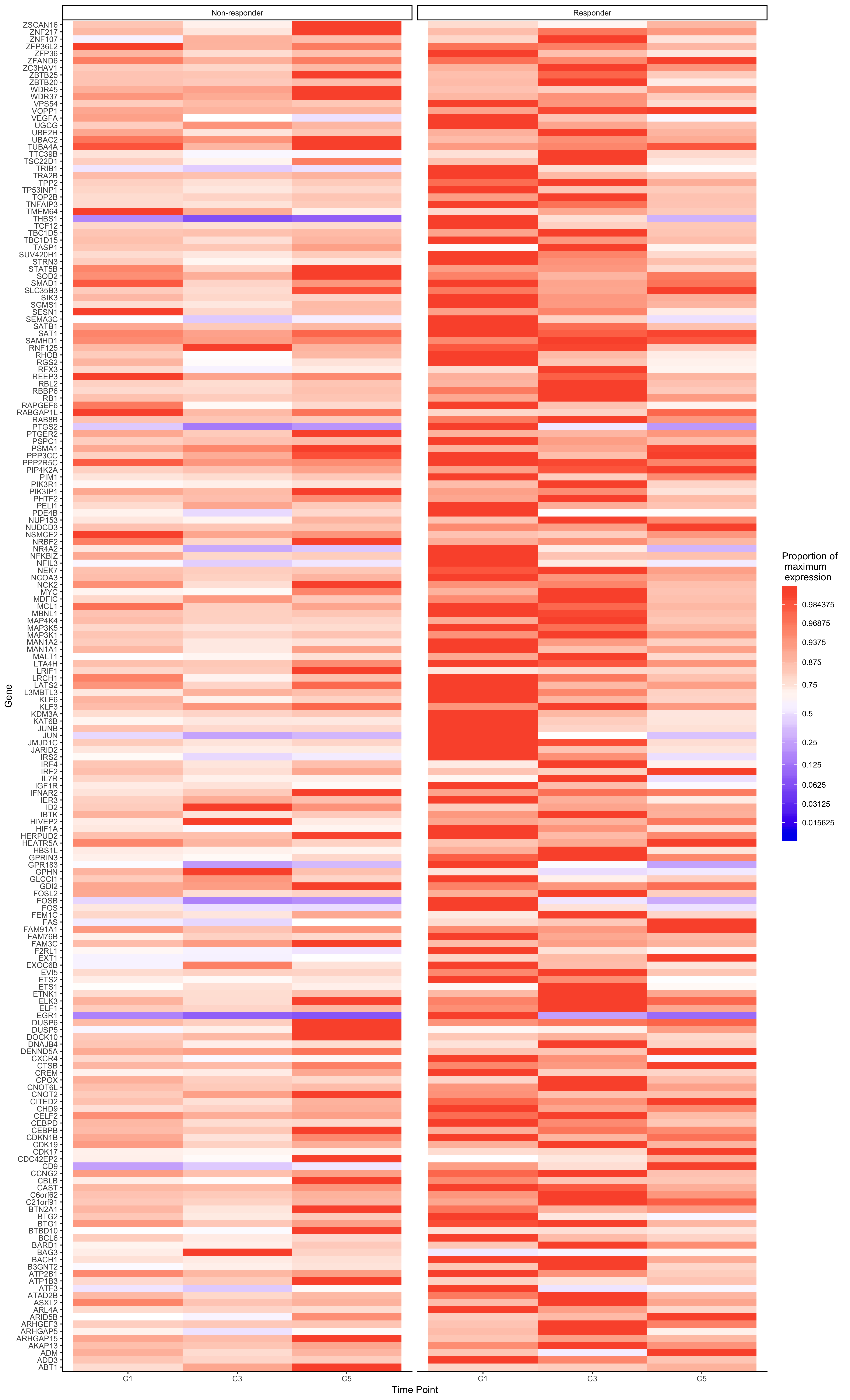

Monocyte differentiation

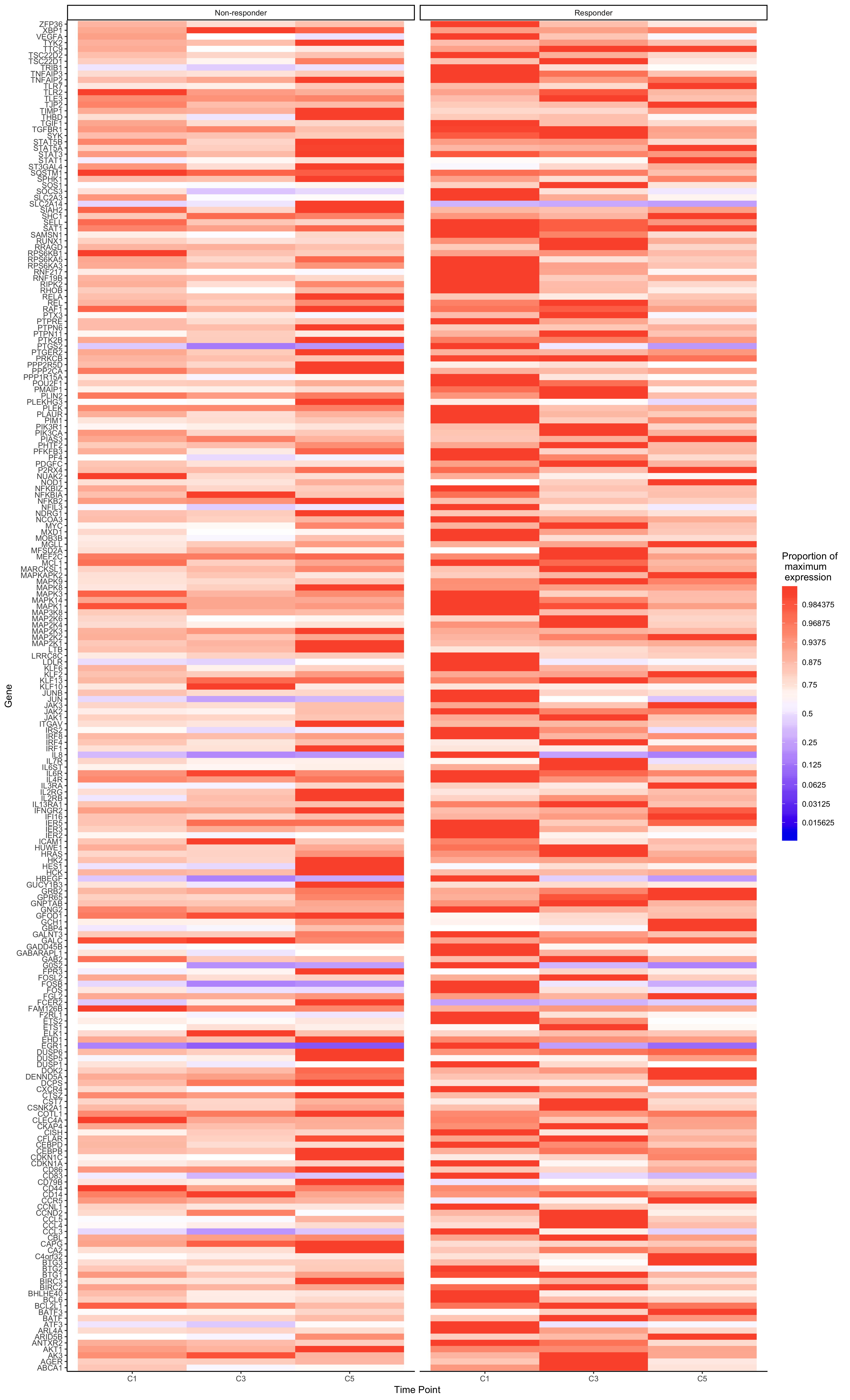

Monocyte inflammation

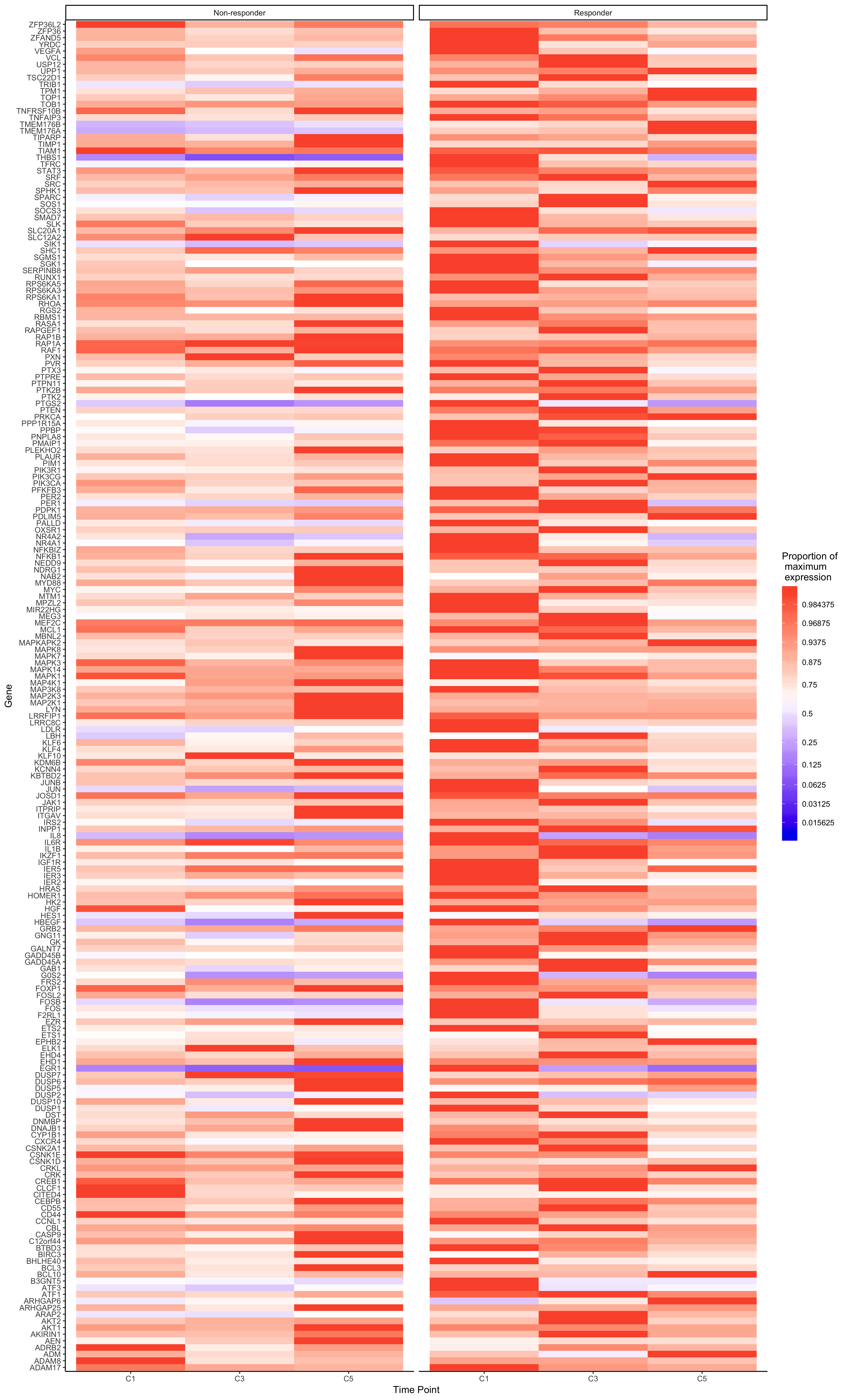

Monocyte growth factors

Fig. S9: Non-responder and responder monocytes show a different gene expression pattern of inflammatory, growth factor, and differentiation pathways before and in response to anti-PD-1 immunotherapy. Heatmaps of changes in monocyte differentiation, growth factor production and inflammatory response gene expression over time in responders and non-responders. Gene expression is displayed as the proportion of the maximum level of each gene. Expression compared at time points C1 (baseline), C3 (start of chemotherapy mFOLFOX6 regimen) and C5 (Two cycles of anti-PD-1 immunotherapy in addition to chemotherapy).

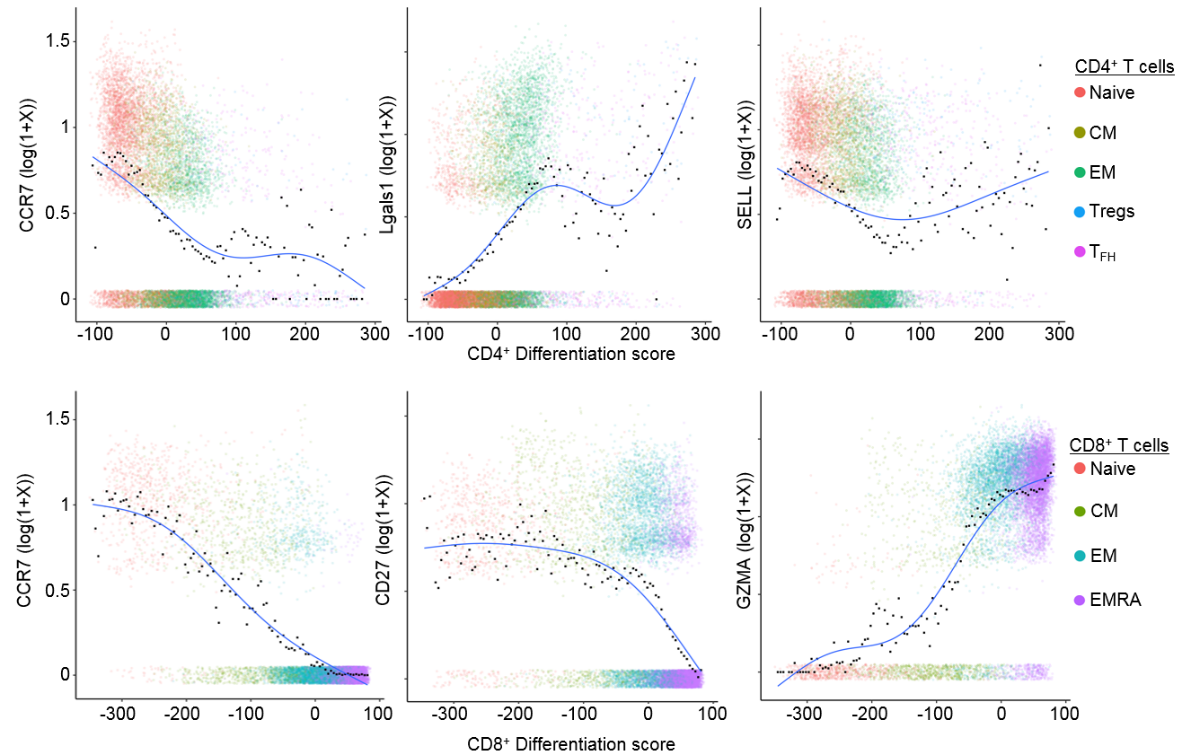

Fig. S10: Phenotype axes capture the continuous spectrum of T cell differentiation states. Changes in CD4^+^ and CD8^+^ T cell gene expression are shown across the T-cell differentiation gradient. For each cell (dot) in each T cell subtype (color), the cell differentiation score is plotted against marker genes of naïve (CCR7) and effector states (LGALS1 and GZMA). Average expression of cells in small regions of the phenotype gradient are calculated, after discretizing the differentiation axis into 100 categories and classifying cells into these groups. A generalized additive model indicates the trend in gene expression across the phenotype gradient.

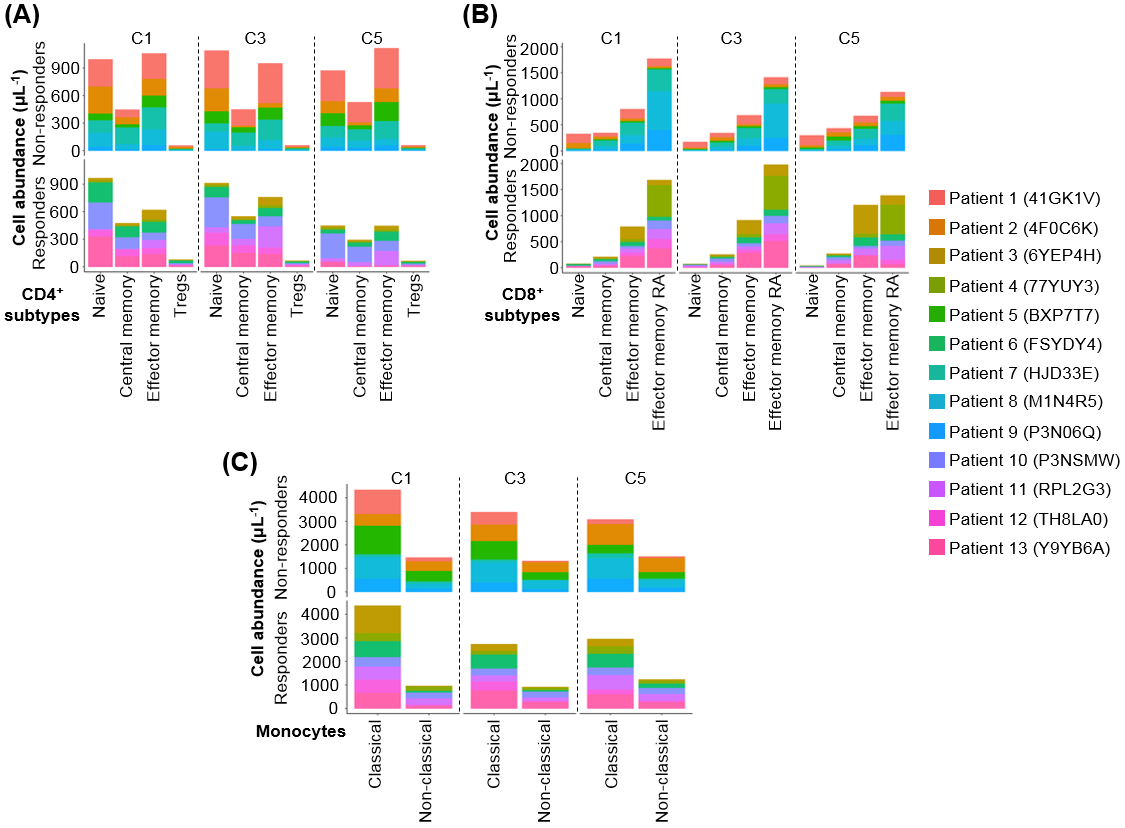

Fig. S11: Abundance of different immune cell subtypes in CD4^+^/CD8^+^ T cells and monocytes differs between responders and non-responders and over time. Immune cell abundances are shown for each patient (color) for subtypes of: **(A)** CD4^+^ T cell, **(B)** CD8^+^ T cell and **(C)** monocytes. Patients are grouped into responders and non-responders in each panel. C stands for cycle, 1 cycle is 14 days, C1= baseline, C3= chemotherapy mFOLFOX6 regimen, C5 and 8= Chemotherapy + anti PD-1 immunotherapy.

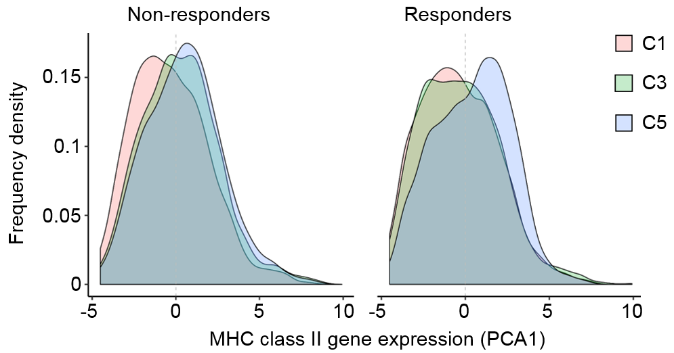

Fig. S12: Major Histocompatibility Complex gene expression of monocytes increases more in responders than non-responders following anti-PD-1 immunotherapy. Frequency of monocytes with different levels of MHC II gene expression compared between responders and non-responders at each treatment time point. The overall MHC II expression score was produced using the first dimension of a principle component analysis applied to the expression of all HLA class II alpha chain genes within each cell.

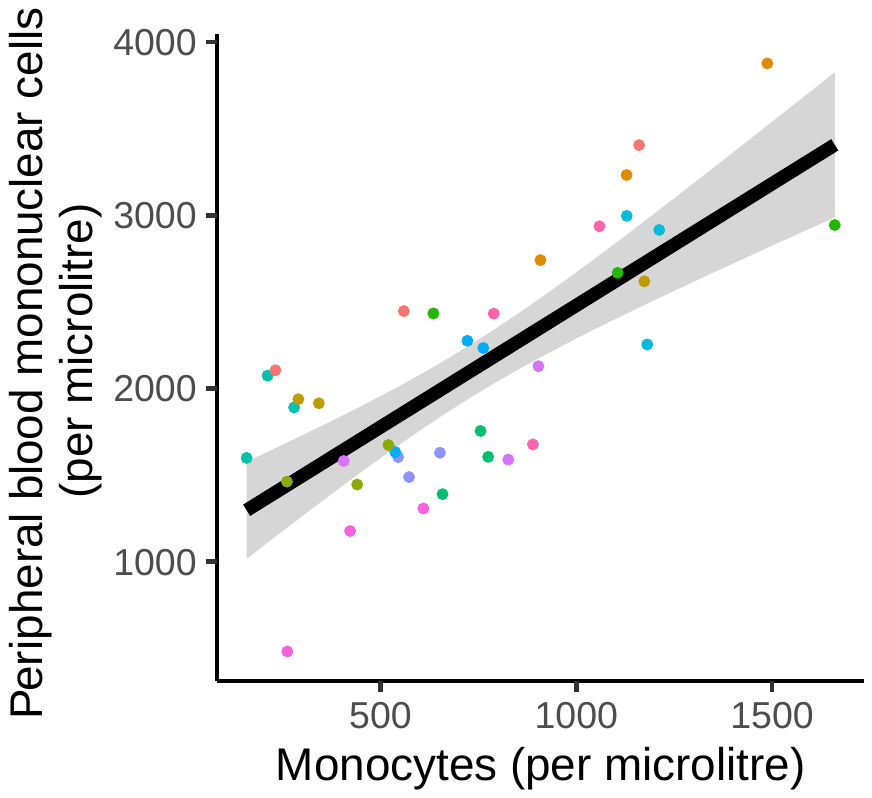

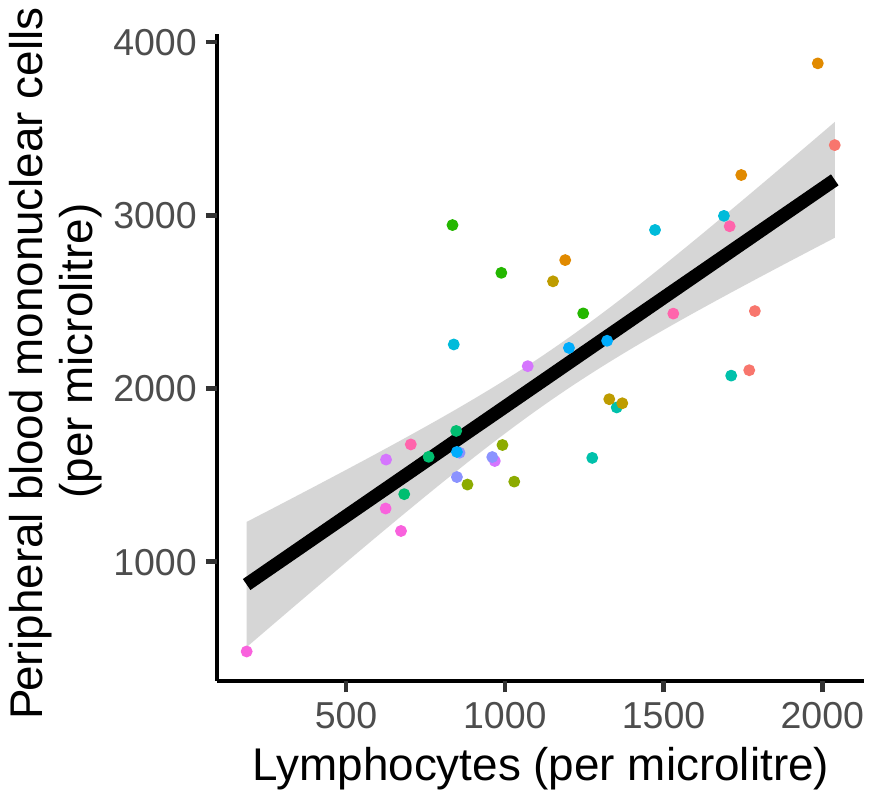

Fig S13: Peripheral blood mononuclear cell abundance of patient’s during the study is positively correlated with Lymphocyte and Monocyte abundance.

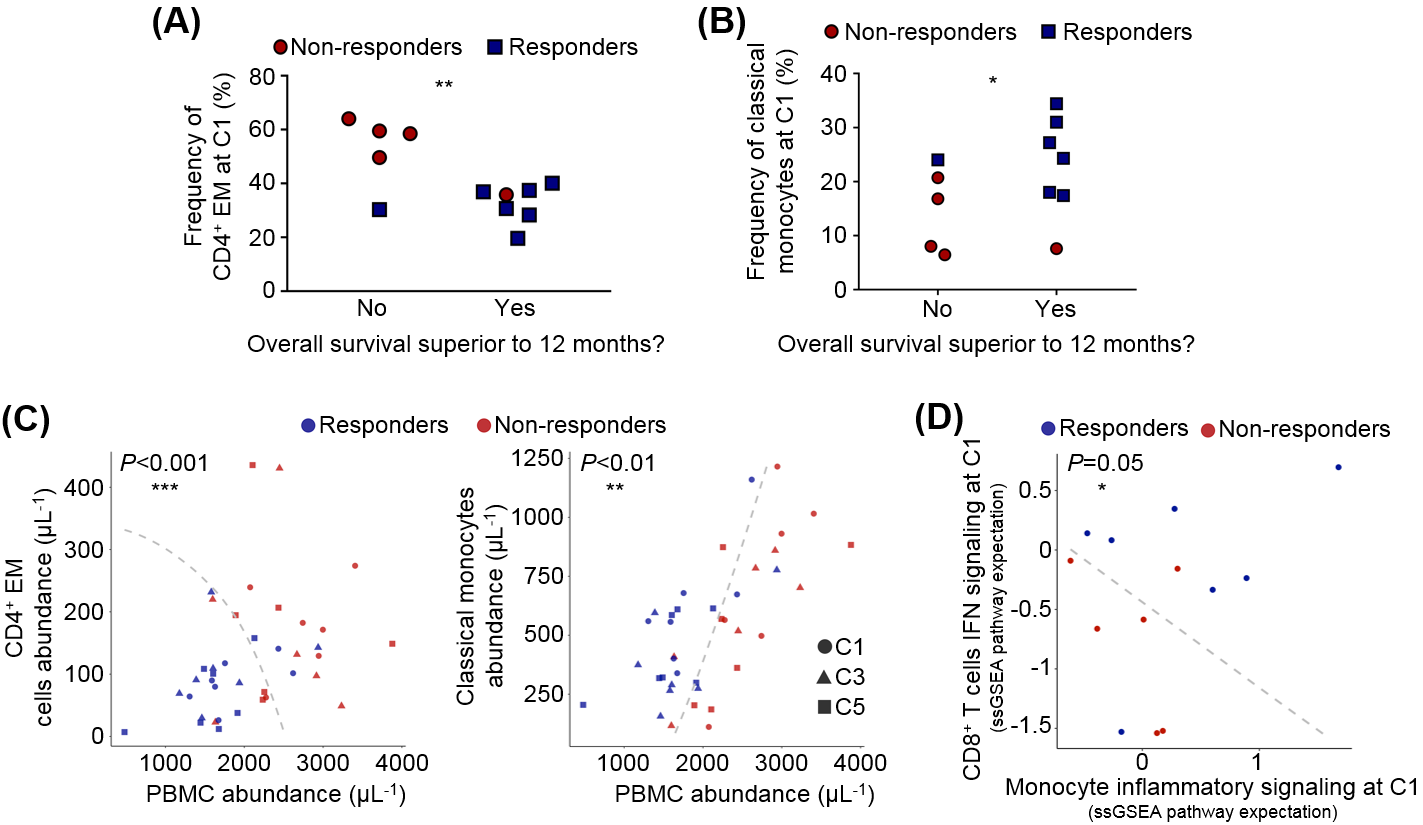
 Fig S14: Abundance and signaling states of CD4^+^ EM cells and monocytes are predictive markers of response to therapy. (**A**) The relationship between CD4^+^ EM frequency observed by flow cytometry and the overall survival at 12 months in non-responders and responders. (**B**) The relationship between classical monocyte frequency observed by flow cytometry and the overall survival at 12 months in non-responders and responders. (**A-B**) ‘'No’' indicates patients with survival less than 12 months while ‘‘Yes’' indicates patients with survival greater than 12 months. (**C**) Response of GI cancer patients to PD-1 inhibitor immunotherapy (point color) depends on PMBC abundance at the early stages of treatment (C1-C5), as well as CD4^+^ EM (Effector memory) T cells and classical monocyte abundances. Patient response boundaries (dashed lines) were predicted using a hierarchical logistic regression model and delineate responsive and non-responsive immune states. (**D**) Pretreatment patient averaged GSEA pathway scores for CD8^+^ T cell interferon (IFN) activation and monocyte inflammatory response. The figure shows the correlation of monocytes inflammatory signaling and CD8^+^ IFN signaling and responsiveness to therapy. (**A-B**) Statistical significance is displayed as *P<0.05 and ***P*<0.01 (two-tailed unpaired *t*-test). C stands for cycle, 1 cycle is 14 days, C1= baseline, C3= chemotherapy mFOLOFX6 regimen, C5= Chemotherapy + anti PD-1 immunotherapy.

### *Frequencies of CD4^+^EM and classical monocytes are correlated with overall survival*

We analyzed the relationship between the frequency of CD4^+^EM cells determined by flow cytometry and the overall survival after 12 months of treatment (**Fig. S14A**). We found that 7 patients had an overall survival greater than 12 months, and their frequency of CD4^+^ EM was below 40% at the C1 time point. Among these 7 patients, 6 were responders and 1 a non-responder. In contrast, 5 patients survived less than 12 months. Four were non-responders and showed a frequency of CD4^+^EM greater than 40%. Similarly, classical monocyte counts prior to therapy also showed a trend toward response to immunotherapy (**Fig. S14B**).

In order to test whether immune cell abundance and signaling states predict response to therapy, we developed random effects logistic regression models. Patients with a lower total PBMC abundance (cells $\mu L^{-1}$) during the first five cycles of treatment were significantly more likely to respond to anti-PD-1 (z=-3.01, p<0.001). Incorporating the frequencies of CD4^+^ T cell and classical monocyte immune sub-classes significantly improved the prediction of patient response (**Fig. S14C**). Patients with a lower abundance of CD4^+^ effector memory cells were significantly more likely to respond to anti-PD-1 therapy than would be expected given their PBMC abundance (z=-4.03, p<0.001). Patients with a high frequency of classical monocytes were also significantly more likely to respond (z=2.76, p<0.01). By combining data from initial PBMC count and immune subclass frequencies, patient responses could be predicted with an overall 85% accuracy (**Fig. S14C**). We next assessed differences in pathway activity signatures (GSEA scores) between responders and non-responders at baseline. Responsive patients had either: a) monocytes exhibiting a high inflammatory response or b) CD8^+^ T cells with high IFN signaling activation (z=3.18, p<0.001) (**Fig. S14D**). These results indicate that patient anti-PD-1 response could be predicted at early stages of therapy based on a low CD4^+^EM T cell count and a high fraction of classical monocytes. Importantly, the signaling state of T cells and monocytes may contribute to outcomes.

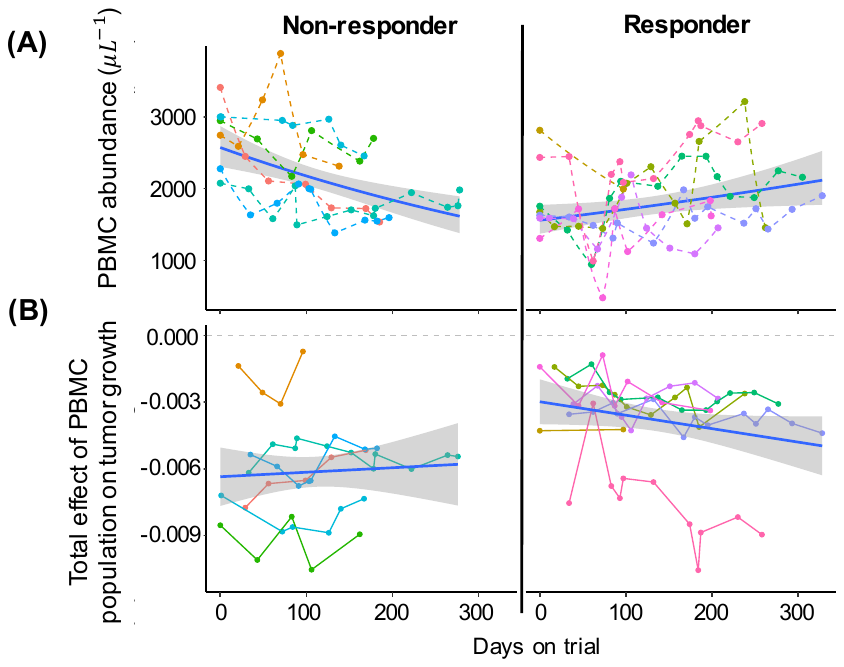

Figure S15: Responsive and non-responsive patients differed in: A) Immune cell abundance dynamics (colored dotted lines=individual responses, Thick blue line=average patient response), and B) the impacts of the immune populations on tumor growth (colored dotted lines= individual model predicted immune effects, Thick blue line=average patient model prediction).

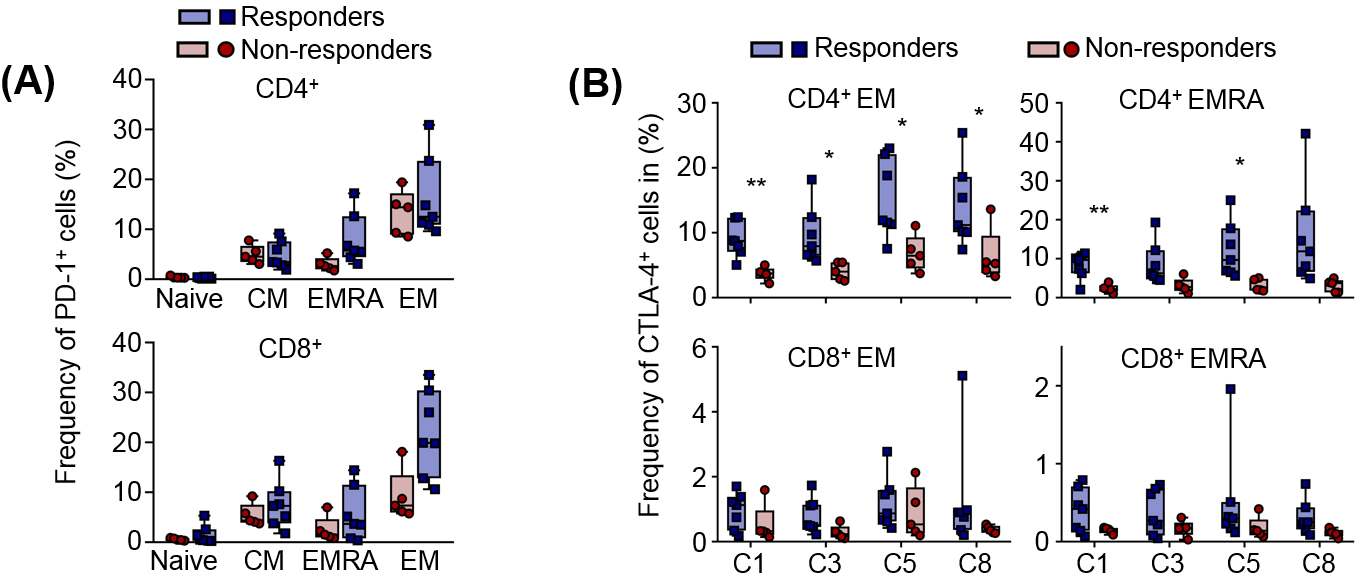

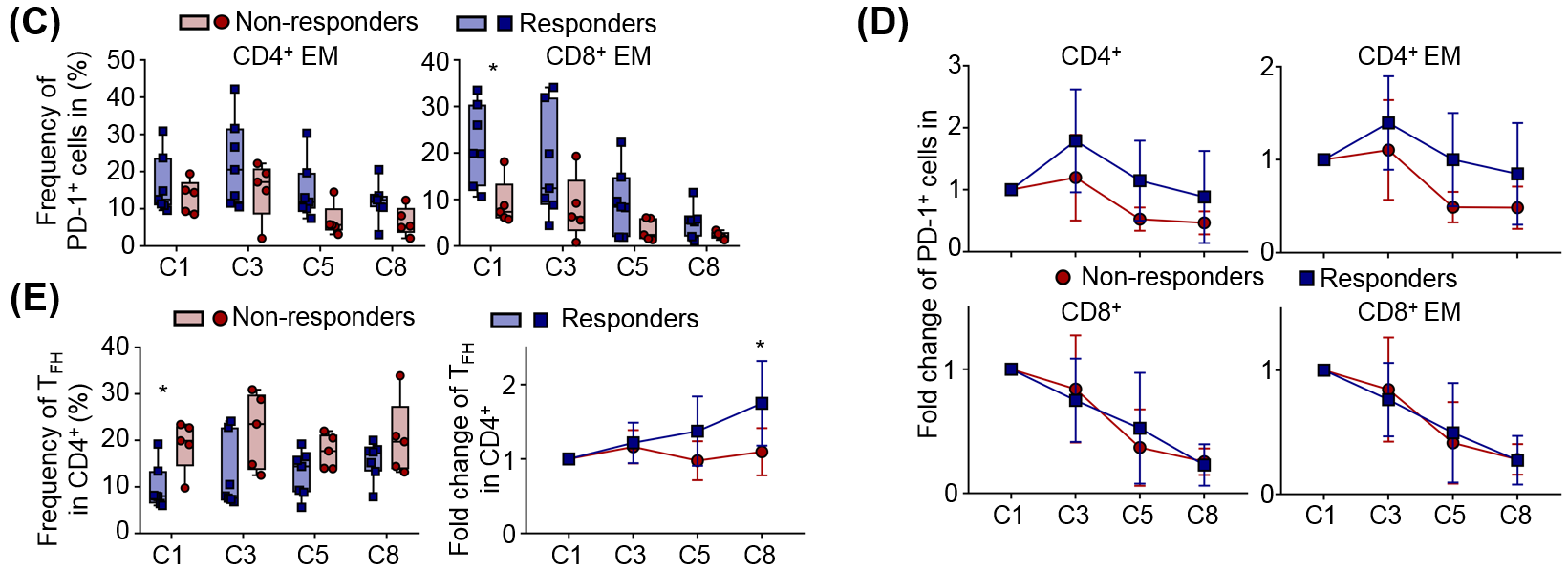

Fig S16: Exhaustion of circulating memory T cells is a hallmark of response prior to therapy. **(A)** Frequencies of PD-1^+^ cells in CD8^+^ (left) CD4^+^ (right) subpopulations at C1. **(B)** Top: frequencies of CTLA4^+^ cells of CD4 EM or EMRA cells and bottom: CD8^+^ EM or EMRA cells. **(C)** Frequencies of PD-1^+^ cells in CD4^+^ EM or CD8^+^ EM cells between responders and non-responders over the course of treatment. **(D)** Fold change of PD1^+^ cells of responders and non-responders over the course of treatment. **(E)** frequencies (left) and fold change (right) of T_FH_ (Follicular helper) between responders and non-responders across treatment. (**A-C**, **E**) Box plots represent the interquartile range (IQR), with the horizontal line indicating the median. Whiskers extend to the farthest data point within a maximum of 1.5× IQR. (**D-E**) Vertical bars indicate s.d. Statistical significance is displayed as *P<0.05, **P<0.01 and ***P<0.001 (two-tailed unpaired *t*-test). C stands for cycle, 1 cycle is 14 days, C1= baseline, C3= chemotherapy mFOLOFX6 regimen, C5, and 8= Chemotherapy + anti-PD-1 immunotherapy.

### *The frequency of PD1*^+^ *exhausted effector T cells decreases during immunotherapy, but the frequency of CTLA-4 exhaustion in these cells increases.*

As pembrolizumab targets and blocks the PD-1 receptor, we evaluated the frequencies of PD-1^+^ cells among CD4^+^ and CD8^+^ T cells. Before treatment, EM cells were the most frequent PD1^+^ subset for both CD4^+^ and CD8^+^ cells (**Fig. S16A**). With the alternative exhaustion marker, CTLA-4, the frequency of CTLA-4^+^ cells within the CD4^+^EM cells and CD4^+^ EMRA populations was higher in responders across all time points (**Fig. S16B**). We found no significant difference in CTLA-4^+^ frequency among CD8^+^ subtypes; however, PD-1^+^ CD8^+^ EM cells were significantly more frequent in responders than non-responders before treatment (**Fig. S16C**).

Across all treatment time points, PD-1 expression in CD8 and CD4 T cells differed between responders and non-responders. In both groups, PD-1^+^ CD8^+^ and PD-1^+^ CD8^+^EM cell frequencies significantly decrease upon commencement of anti-PD-1 therapy. CD4^+^ and CD4^+^EM cell frequencies showed a significant decrease only in the non-responder patients after the start of anti-PD-1 (*p*<0.05) (**Fig. S16D**). This decrease of PD-1 levels by FACS may be explained by the internalization of the receptor after binding pembrolizumab, suggesting that PD-1^+^ CD8^+^ T cells, CD4^+^, and especially CD4^+^EM cells of non-responders are targeted by the drug. CD4^+^ EM cells expressed PD-1 and were found at a higher frequency in non-responders, suggesting that these cells may act as a collateral target of pembrolizumab reducing its availability to cytotoxic T cells. In addition, the frequency of PD-1^+^ T follicular helper cells (T_FH_), a population known to highly express PD-1 (39), was higher in non-responders than responders at C1. (**Fig. S16E**).

SUPPLEMENTARY tables

|  | **All Patients (n=17)** | **Responders (n=8)** | **Non-responders (n=9)** |
| --- | --- | --- | --- |
| **Age, median years (IQR)** | 63 (55-71) | 67 (60-73) | 60 (48-66) |
| **Gender, n (%)** |  |  |  |
| F | 7 (41) | 2 (25) | 5 (56) |
| M | 10 (59) | 6 (75) | 4 (44) |
| **Race, n (%)** |  |  |  |
| White | 13 (76) | 8 (100) | 5 (56) |
| Black or African American | 2 (12) | 0 (0) | 2 (22) |
| Unknown | 2 (12) | 0 (0) | 2 (22) |
| **ECOG Performance Status, n (%)** |  |  |  |
| 0 Fully active | 10 (59) | 4 (50) | 6 (67) |
| 1 Restricted | 7 (41) | 4 (50) | 3 (33) |
| **Primary Cancer Type, n (%)** |  |  |  |
| Colorectal Carcinoma | 8 (47) | 3 (38) | 5 (56) |
| Biliary Tract Cancer | 4 (24) | 1 (13) | 3 (33) |
| Gastroesophageal Carcinoma | 3 (18) | 3 (38) | 0 (0) |
| Pancreatic Carcinoma | 2 (12) | 1 (13) | 1 (11) |
| **Pembrolizumab Dose level, n (%)** |  |  |  |
| 75 mg q 2 weeks | 1 (6) | 0 (0) | 1 (11) |
| 200 mg q 2 weeks | 16 (94) | 8 (100) | 8 (89) |
| **Prior Systemic Treatment, n (%)** |  |  |  |
| No | 6 (35) | 6 (75) | 0 (0) |
| Yes | 11 (65) | 2 (25) | 9 (100) |
| Days from last dose of treatment, Median (IQR) | 41 (34-167) | 101 (34-167) | 41 (31-361) |
| **Prior 5-FU and Oxaliplatin, n (%)** |  |  |  |
| No | 10 (59) | 7 (88) | 3 (33) |
| Yes | 7 (41) | 1 (13) | 6 (67) |
| Days from last dose of 5-FU/Ox, Median (IQR) | 657 (249-1005) | 927 (927-927) | 552 (220-1314) |
| **Number of previous systemic treatment lines for metastatic disease, n (%)** |  |  |  |
| 0 | 7 (41) | 6 (75) | 1 (11) |
| 1 | 4 (24) | 2 (25) | 2 (22) |
| 2 | 3 (18) | 0 (0) | 3 (33) |
| 3 | 3 (18) | 0 (0) | 3 (33) |
| **Microsatellite Instability (MSI) Result, n (%)** |  |  |  |
| dMMR (MSI-high) | 4 (24) | 3 (38) | 1 (11) |
| MMR proficient (MSI-stable) | 7 (41) | 1 (13) | 6 (67) |
| Unknown | 6 (35) | 4 (50) | 2 (22) |
| **Correlative testing, n (%)** |  |  |  |
| FACS | 4 (24) | 1 (13) | 3 (33) |
| Single cell (SC) | 5 (29) | 1 (13) | 4 (44) |
| FACS and SC | 8 (47) | 6 (75) | 2 (22) |

Table S1: Patient and disease characteristics

| **Blood Count with Differential, laboratory reference range** | **All Patients**  **(n = 17)** | | **Responders**  **(n=8)** | | **Non-responders (n=9)** | |
| --- | --- | --- | --- | --- | --- | --- |
|  | **Median (IQR)** | **Range** | **Median (IQR)** | **Range** | **Median (IQR)** | **Range** |
| **White Blood Cells, 3.20 - 10.60 k/μL** |  |  |  |  |  |  |
| C1 | 6.8 (3.9) | 3.9-11.4 | 7.3 (5.4) | 4.5-11.4 | 5.9 (3.6) | 3.9-10.4 |
| C3 | 4.1 (2.3) | 2.5-6.7 | 4.5 (1.4) | 3.1-6.7 | 3.5 (2.6) | 2.5-6.7 |
| C5 | 4.3 (1.9) | 2.4-11.9 | 4.7 (1.7) | 3.2-10.5 | 3.9 (2.1) | 2.4-11.9 |
| **Red Blood Cells, 3.88 - 5.46 M/ μL** |  |  |  |  |  |  |
| C1 | 4.4 (0.7) | 3.5-5.4 | 4.7 (0.6) | 3.8-5.4 | 4.1 (0.8) | 3.5-5.0 |
| C3 | 4.1 (0.9) | 2.8-5.4 | 4.4 (0.9) | 3.1-5.4 | 3.9 (0.8) | 2.8-4.8 |
| C5 | 4.0 (1.2) | 2.8-5.3 | 4.4 (0.9) | 2.8-5.3 | 3.6 (1.0) | 3.4-4.6 |
| **Hemoglobin, 12.1 - 15.9 g/dL** |  |  |  |  |  |  |
| C1 | 12.8 (2.5) | 9.4-16.1 | 12.8 (2.5) | 9.4-15.2 | 12.9 (2.9) | 10.0-16.1 |
| C3 | 12.1 (2.1) | 9.5-15.2 | 11.7 (3.3) | 9.5-14.7 | 12.3 (2.9) | 9.7-15.2 |
| C5 | 11.8 (2.6) | 8.8-15.9 | 12.3 (2.4) | 8.9-14.7 | 11.4 (3.5) | 8.8-15.9 |
| **Platelets, 177 - 406 k/ μL** |  |  |  |  |  |  |
| C1 | 204 (78) | 129-353 | 195 (104) | 148-339 | 219 (91) | 129-353 |
| C3 | 147 (53) | 93-228 | 134 (76) | 93-223 | 167 (45) | 112-228 |
| C5 | 139 (70) | 77-404 | 114 (119) | 77-404 | 155 (51) | 120-226 |
| **Granulocyte #,  1.3 - 7.0 k/ μL** |  |  |  |  |  |  |
| C1 | 4.5 (3.2) | 2.6-8.6 | 5.4 (4.5) | 3.0-8.6 | 3.9 (2.0) | 2.6-7.2 |
| C3 | 2.5 (1.8) | 1.1-4.8 | 2.5 (1.1) | 2.0-4.8 | 1.6 (1.8) | 1.1-4.2 |
| C5 | 3.0 (2.1) | 1.1-9.6 | 2.7 (1.4) | 1.4-8.5 | 3.2 (2.6) | 1.1-9.6 |
| **Monocyte #,   0.2 - 0.7 k/ μL** |  |  |  |  |  |  |
| C1 | 0.4 (0.3) | 0.2-1.1 | 0.5 (0.5) | 0.3-1.1 | 0.4 (0.4) | 0.2-0.9 |
| C3 | 0.4 (0.3) | 0.2-0.9 | 0.4 (0.5) | 0.3-0.9 | 0.3 (0.3) | 0.2-0.6 |
| C5 | 0.4 (0.3) | 0.0-1.1 | 0.5 (0.6) | 0.0-1.1 | 0.4 (0.2) | 0.3-0.7 |
| **Lymphocyte #, 0.8 - 3.1 k/ μL** |  |  |  |  |  |  |
| C1 | 1.6 (1.0) | 0.4-2.8 | 1.1 (0.7) | 0.4-1.7 | 1.7 (1.2) | 0.7-2.8 |
| C3 | 1.3 (0.7) | 0.5-2.3 | 1.1 (0.4) | 0.5-1.6 | 1.6 (0.8) | 0.7-2.3 |
| C5 | 1.3 (0.9) | 0.4-3.0 | 1.2 (0.4) | 0.4-2.2 | 1.6 (0.9) | 0.8-3.0 |

Table S2: Blood count parameters on day 1 of cycle 1, 3, and 5.

| Cluster | Major clusters | Sub clusters |
| --- | --- | --- |
| 1 | NK Cell | NK Cell Resting |
| 2 | Monocyte | Monocyte Classical |
| 3 | Monocyte | Monocyte Classical |
| 4 | Monocyte | Monocyte Non-Classical |
| 9 | Monocyte | Monocyte Classical Platelets |
| 10 | Monocyte | Monocyte Classical |
| 11 | B Cell | B Cell Naive |
| 12 | NK Cell | NK Cell Activated |
| 13 | B Cell | B Cell Memory |
| 14 | Dendritic Cell | Dendritic Cell |
| 15 | Monocyte | Monocyte Classical |
| 17 | Cell Cycle | Cell Cycle |
| 18 | Monocyte | Monocyte Classical |
| 19 | Plasmacytoid dendritic cell | Plasmacytoid Dendritic Cell |
| 20 | Plasma Cell | Plasma Cell |
| 22 | Activated Platelets | Activated Platelets |
| 23 | T Cell | NA |
| 24 | RBC | RBC |
| 25 | Plasma Cell | Plasma Cell |
| T0 | T Cell CD4 | T Cell CD4 EM |
| T1 | T Cell CD4 | T Cell CD4 Naive |
| T10 | T Cell CD8 | T Cell CD8 Exhausted |
| T11 | T Cell CD8 | T Cell CD8 EM |
| T12 | T Cell CD8 | T Cell CD8 EM |
| T13 | T Cell CD8 | T Cell CD8 EM |
| T14 | T Cell CD8 | T Cell CD8 CM |
| T15 | T Cell CD4 | T Cell T_FH_ |
| T2 | T Cell CD8 | T Cell CD8 TEMRA |
| T3 | T Cell CD8 | T Cell CD8 TEMRA |
| T4 | T Cell CD4 | T Cell CD4 CM |
| T5 | T Cell CD8 | T Cell CD8 EM |
| T6 | NA | NA |
| T7 | T Cell CD8 | T Cell CD8 CM |
| T8 | T Cell CD4 | T Cell Treg |
| T9 | T Cell CD8 | T Cell CD8 Naive |

Table S3: Major clusters and sub clusters annotation of the UMAP.

**Other supplementary data for this manuscript**

Dataset S1: Clinical data used to parameterize the tumor-immune interaction mathematical model.

Dataset S2: P-values of differential gene expression between time point of patients.

Dataset S3: Significant ssGSEA pathways activation per immune cell subtypes in responders and non-responders.

Dataset S4: Abundance of total PBMC per patients and time points.

**References**

1.	E. Azizi *et al.*, Single-Cell Map of Diverse Immune Phenotypes in the Breast Tumor Microenvironment. *Cell* **174**, 1293-1308 e1236 (2018).

2.	M. Sade-Feldman *et al.*, Defining T-cell States Associated with Response to Checkpoint Immunotherapy in Melanoma. *Cell* **175**, 998-1013 e1020 (2018).

3.	A. Butler, P. Hoffman, P. Smibert, E. Papalexi, R. Satija, Integrating single-cell transcriptomic data across different conditions, technologies, and species. *Nat Biotechnol* **36**, 411-420 (2018).

4.	B. Bischl *et al.*, mlr: Machine Learning in R. *JMLR* **17**, 1-5 (2016).

5.	K. Moon *et al.*, Manifold Learning-based Methods for Analyzing Single-Cell RNA-Sequencing Data. *Current Opinion in Systems Biology* **7**, (2017).

6.	D. van Dijk *et al.*, Recovering Gene Interactions from Single-Cell Data Using Data Diffusion. *Cell* **174**, 716-729 e727 (2018).

7.	K. Moon *et al.*, Visualizing Structure and Transitions for Biological Data Exploration. *SSRN Electronic Journal*, (2018).
